# Core circadian clock and light signaling genes brought into genetic linkage across the green lineage

**DOI:** 10.1101/2021.11.02.466975

**Authors:** Todd P. Michael

## Abstract

The circadian clock ensures that biological processes are phased to the correct time of day. In plants the circadian clock is conserved at both the level of transcriptional networks as well as core genes. In the model plant *Arabidopsis thaliana,* the core circadian *singleMYB* (*sMYB*) genes *CCA1* and *RVE4* are in genetic linkage with the *PSEUDO-RESPONSE REGULATOR* (*PRR*) genes *PRR9* and *PRR7* respectively. Leveraging chromosome-resolved plant genomes and syntenic ortholog analysis it was possible to trace this genetic linkage back to the basal angiosperm *Amborella* and identify an additional evolutionarily conserved genetic linkage between *PIF3* and *PHYA*. The *LHY/CCA1-PRR5/9, RVE4/8-PRR3/7* and *PIF3-PHYA* genetic linkages emerged in the bryophyte lineage and progressively moved within several genes of each other across an array of higher plant families representing distinct whole genome duplication and fractionation events. Soybean maintains all but two genetic linkages, and expression analysis revealed the *PIF3-PHYA* linkage overlapping with the E4 maturity group locus was the only pair to robustly cycle with an evening phase in contrast to the *sMYB-PRR* morning and midday phase. While most monocots maintain the genetic linkages, they have been lost in the economically important grasses (Poaceae) such as maize where the genes have been fractionated to separate chromosomes and presence/absence variation results in the segregation of *PRR7* paralogs across heterotic groups. The evolutionary conservation of the genetic linkage as well as its loss in the grasses provides new insight in the plant circadian clock, which has been a critical target of breeding and domestication.

**Summary Sentence:** The genetic linkage of the core circadian clock components has evolutionary origins in bryophytes and sheds light on the current functioning and selection on the circadian clock in crops.

## Introduction

The plant circadian clock ensures biological processes occur at the proper time-of-day (TOD) regardless of predictable as well as random environmental changes, which is why it has been a target for plant domestication (Bendix et al., 2015; Creux and Harmer, 2019; McClung, 2021; Steed et al., 2021). The molecular mechanisms of the plant circadian clock were initially worked out in the model plant *Arabidopsis thaliana* and now there is a growing body of work across an array of crop and ornamental plants (Sanchez and Kay, 2016; McClung, 2019). The circadian clock is so named because the period is approximately a day (*circa diem*) varying between 22 and 27 hrs across different *Arabidopsis* accessions, which correlates with the latitude of origin (Michael et al., 2003). The almost 24 hr period of the circadian clock enables plants to anticipate changes in photoperiod each day over the growing season synchronizing timing of biological processes and ultimately enhancing plant fitness (Green et al., 2002; Dodd et al., 2005). Fundamental processes in plants are under the regulation of the circadian clock such as phytohormone-regulated growth, photosynthetic gene expression, and the cell cycle (Michael et al., 2008a; Fung-Uceda et al., 2018). While the specific molecular structure of the clock may vary across the green kingdom, many features including gene content and expression patterns are conserved (Ferrari et al., 2019; Wickell et al., 2021).

Autotrophic organisms that rely on photosynthesis for energy from algae to plants are highly dependent on synchronizing their biology to the daily changes in light and temperature. For instance, almost all of the genes in the single celled pico-algae *Ostreococcus* show oscillations in gene expression in a TOD fashion (Monnier et al., 2010; Thommen et al., 2012), while 80% of genes in the model macroalga *Chlamydomonas reinhardtii* have TOD peak abundance (Zones et al., 2015). In most higher plants 30-40% of genes cycle with cycling of light and temperature conditions, and 10-20% cycle under continuous light and temperature (Michael et al., 2008b; Khan et al., 2010; Filichkin et al., 2011; Sato et al., 2013; Nose and Watanabe, 2014; Cronn et al., 2017; Ferrari et al., 2019; MacKinnon et al., 2019; Wai et al., 2019; Greenham et al., 2020; Lai et al., 2020; Michael et al., 2020; Wickell et al., 2021). However, there is evidence that the majority of genes in higher plants may have TOD expression potential since in *Arabidopsis* 90% of all genes have TOD expression in at least one of an array of conditions tested (Michael et al., 2008b). Now there is an emerging body of literature concerning the circadian clocks across an array of plants and how they respond to natural conditions of light and temperature (Panter et al., 2019).

The molecular architecture and genes involved with the plant circadian clock are best studied in *Arabidopsis*, which has revealed a highly complex network of negative and positive feedback loops with about 61 genes (Lou et al., 2012; McClung, 2019). At the core of these feedback loops are two gene families: the *single MYB* (*sMYB*) transcription factors *LATE ELONGATED HYPOCOTYL* (*LHY*), *CIRCADIAN CLOCK ASSOCIATED 1* (*CCA1*) and *REVEILLE* (*RVE*) family, and the *PSEUDO RESPONSE REGULATOR* (*PRR*) family (McClung, 2019). The *RVE* family is composed of 8 members: *RVE1*, *RVE2*, *RVE3*, *RVE4*, *RVE5*, *RVE6*, *RVE7* and *RVE8*; all expressed in the morning or midday (Supplemental Figure 1) (Michael et al., 2008b; Rawat et al., 2009; Gray et al., 2017). The *PRR* family is comprised of 5 members: *PRR1*, *PRR3*, *PRR5*, *PRR7*, and *PRR9*, which have peak expression over the entire day starting at dawn (*PRR9*), midday (*PRR7*), evening (*PRR3* and *PRR5*) and early night (*PRR1* commonly called *TOC1*) (Supplemental Figure 1) (Matsushika et al., 2000; Michael et al., 2008b). A highly simplified model of these negative and positive feedback loops has *CCA1/LHY* negatively regulating the *PRRs*, which in turn negatively regulate both *CCA1/LHY* and *RVEs*, while the *RVEs* play a positive role in positively regulating the *PRRs* (Figure 1A) (Shalit-Kaneh et al., 2018).

The circadian clock is also conserved from algae to higher plants at both the level of the proteins as well as TOD networks (Filichkin et al., 2011; Reyes et al., 2017). While higher plants have more components, this core negative feedback circadian clock is conserved as far back as *Ostreococcus,* whose oscillator is made up of one *sMYB-PRR* loop (Corellou et al., 2009; Monnier et al., 2010; Morant et al., 2010). Subsequently, several studies have explored the evolution and conservation of the core circadian clock components in bryophytes, lycophytes and a diverse array of angiosperms (Holm et al., 2010; McClung, 2010; Takata et al., 2010; Satbhai et al., 2011; Ryo et al., 2016; Linde et al., 2017; Wickell et al., 2021). It was found that core clock genes are preferentially retained after the triplication in *Brassica rapa*, consistent with the augmented roll of the circadian clock during domestication (Lou et al., 2012; McClung, 2021). While these components are conserved all the way back to algae from higher plants, it is not clear how they have been inherited over evolutionary time.

It was previously noted that *CCA1* and *PRR9* are in genetic linkage on Chromosome 2 (Chr02) in *Arabidopsis* (Michael et al., 2003; Lou et al., 2012), and later that *RVE4* and *PRR7* are linked on Chr05 (Michael et al., 2008b). However, despite the orthology of these *sMYB-PRR* gene pairs, these regions were not found to be syntenic in *Arabidopsis* (Lou et al., 2012). Similar *sMYB-PRR* linkages have been noted in other dicot species such as cranberry and blueberry, and monocot species such as duckweed (*Wolffia*, *Lemna*, *Spirodela*) (Michael et al., 2020; Abramson et al., 2021; Kawash et al., 2021). These results suggest the *sMYB-PRR* linkage is evolutionarily conserved and that it may predate the land plants. Since these pairs are in genetic linkage, double mutants are not yet available to assess the significance. Therefore, a syntenic ortholog analysis was conducted across the green kingdom to elucidate the evolutionary linkage between *sMYB-PRR*.

## Results

### Syntenic paralogs in *Arabidopsis thaliana* clock genes

For the purpose of exploring the relationship between *CCA1-PRR9* and *RVE4-PRR7*, a simplified model of their interactions was developed (Figure 1A) (Shalit-Kaneh et al., 2018); more detailed models can be found in recent reviews (McClung, 2021; Steed et al., 2021). In general, *PRR7/9* are negatively and positively regulated by *CCA1/LHY* and *RVE4/RVE8* respectively, while in turn *PRR7/9* negatively regulate both. Consistent with these reciprocal functions, loss-of-function mutants *cca1* and *lhy* result in short circadian period lengths (<24 hrs), while loss-of-function *prr7*, *prr9*, *prr7/9*, and rve4/6/8 result in long circadian period lengths (>24 hrs) (Figure 1A). *CCA1*, *LHY*, *RVE4*, and *RVE8* have been shown to be expressed at dawn or shortly after; In contrast, *PRR9* has been shown to be expressed 4 hrs after dawn (Zeitgeber 4; ZT4) and *PRR7* at ZT7 (Figure 1B,C; Supplemental Figure S1). Therefore, *PRR9* and *PRR7* peak several hours after their syntenic pairs of *CCA1* and *RVE4 respectively*.

**Figure 1.**
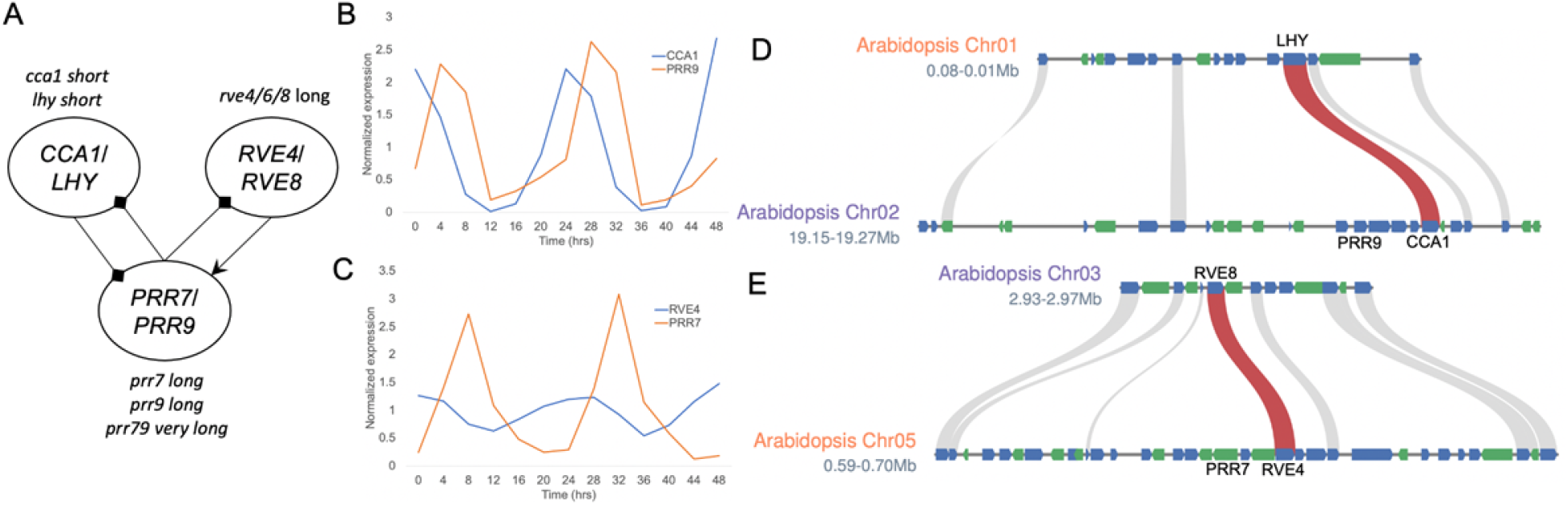
Core circadian clock genes in *Arabidopsis* are found in tight linkage. A) Simplified plant circadian clock model that includes the single MYB (sMYB) paralogs *CCA1*, *LHY*, *RVE4*, and *RVE8*, and the *PSEUDO-RESPONSE REGULATORS* (*PRRs*) *PRR7* and *PRR9*. Paralogs *CCA1/LHY* and *RVE4/8* are expressed in the morning with the former negatively regulating and the latter positively regulating the midday expressed *PRR7/9* paralogs. The circadian phenotype of the knockout mutants is included: long, long period (>24 hrs); short, short period (<24 hrs). B) *CCA1* (blue) and *PRR9* (orange) display morning (Zeitgeber Time 0; ZT0) and midday (ZT4) peak time-of-day (TOD) expression respectively. C) *RVE4* (blue) and *PRR7* (orange) display morning (ZT4) and midday (ZT7) peak time-of-day (TOD) expression respectively. D) Syntenic regions on *Arabidopsis* chromosome 1 (Chr01; orange) containing *LHY* and Chr02 (purple) containing *CCA1*. *PRR9* is found 3 genes away from *CCA1*. E) Syntenic regions on *Arabidopsis* Chr03 (purple) containing *RVE8* and Chr05 (orange) containing *RVE4*. *PRR7* is found 2 genes away from *RVE4*. The other syntenic genes (forward, blue; reverse, green) are depicted in the homologous chromosomal regions (grey).

The tight linkage of both sets of *CCA1-PRR9* and *RVE4-PRR7* paralogs suggests this could be mechanistically important to the daily timing of the clock. Additional linked genes and syntenic blocks were searched for using a collection of 61 circadian clock and light signalling genes (Supplemental Table S1) (Lou et al., 2012). There were five gene pairs that were found within 12 genes of one another: *CCA1-PRR9* (4), *RVE4-PRR7* (3), *PHYA-PIF3* (4), *PHYB-LKP2* (12) and *SRR1-BOA* (1) (gene pair and number of intervening genes) (Supplementary note). In addition, 38% (23) of genes were found in syntenic blocks with 14 in syntenic paralog pairs, and 6 genes in the syntenic pairs not on the circadian clock gene list (Supplemental Table S2). As has been reported before, the core clock genes *CCA1*-*LHY*, *RVE3*-*RVE5*, *ZTL*-*LKP2* are in syntenic blocks (Lou et al., 2012) as well as *RVE8*-*RVE4* (Figure 1D,E; Supplemental Figure 2; Supplemental Table S2) (Shalit-Kaneh et al., 2018). However, no syntenic relationships were identified between the closely linked genes, nor specifically the *CCA1-PRR9* and *RVE4-PRR7* blocks. This result suggests that the *CCA1-PRR9* and *RVE4-PRR7* blocks were differentially inherited, possibly through distinct whole genome duplication (WGD) and fractionation events.

*Arabidopsis* has experienced three WGD events termed lambda (◻), beta (◻) and alpha (◻) (Jiao et al., 2014). Based on Ks (synonymous substitutions) for the syntenic blocks, *CCA1*-*LHY* and *ZTL*-*LKP2* emerged from the ◻ WGD ~150 million years ago (mya); *LUX-BOA*, *RVE3*-*RVE5*, and *CHE-TCP7* emerged from the ◻ ~75 mya; and *RVE4*-*RVE8* and *FLC-MAF5* emerged in the most recent ◻ WGD ~50 mya (Supplemental Table S2). This means that *CCA1-LHY* ortholog copies have been purged during the ◻ and ◻ events to maintain a single copy of each while *RVE4-RVE8* emerged recently, consistent with the two gene families representing two distinct evolutionary trajectories. Consistent with this result, a pair-wise Ks analysis of both the *sMYB* and *PRR* families suggested the paralogs in each family are evolutionarily distant while *RVE4-RVE8* is not since it is the result of the most recent ◻ WGD event (Supplemental Tables S3; Supplemental Table S4). The synteny analysis and the Ks analysis both suggest that the *sMYB-PRR* linkage arose earlier than the *Arabidopsis* lineage.

### sMYB-PRR linkage in basal angiosperm *Amborella*

*Amborella trichopoda* is a basal angiosperm that gave rise to both dicots and monocots and lacks a WGD event, which makes it attractive for tracing the lineage of gene families (Amborella Genome Project, 2013). *Amborella* only has three *PRR* proteins that are orthologous to *Arabidopsis PRR1*, *PRR3/7* and *PRR5/9*; the *PRR5/9* ortholog was split during gene prediction as two genes (ATR0661G529_ATR0661G570) (Figure 2A). In addition, *Amborella* has four *sMYB* genes that are orthologs of *Arabidopsis CCA1/LHY*, *RVE4/8*, *RVE1/2/7*, and *RVE6* consistent with the fact that the ancestral plant has both *CCA1/LHY* and *RVE4/8* orthologs (Figure 2B) (Sharma et al., 2017; Shalit-Kaneh et al., 2018). *Amborella* does not have a *RVE3/5* pair, which could mean that the pair arose from *RVE6* as suggested (Lou et al., 2012).

**Figure 2.**
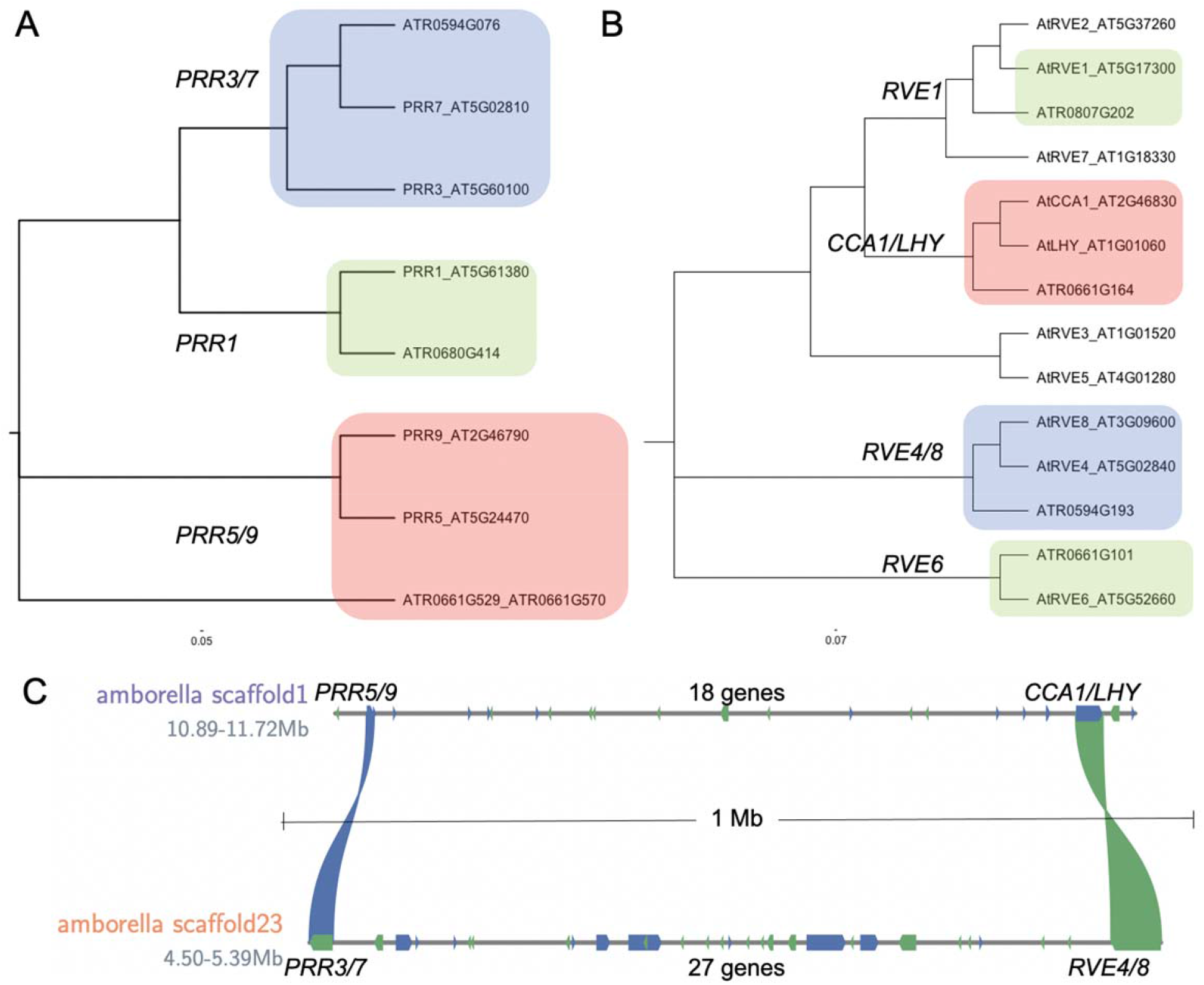
The basal plant *Amborella trichopoda* has *sMYB-PRR* associations. The *Amborella* (*Amborella trichopoda*) and *Arabidopsis* (*Arabidopsis thaliana*) A) *PRR* and B) *sMYB* phylogenetic trees resolve the specific relationships. C) *Amborella* scaffold01 and scaffold23 have the *CCA1/LHY-PRR5/9* and *RVE4/8-PRR3/7* linkages 18 and 27 genes apart respectively. Other genes (forward, blue; reverse, green) are depicted on the scaffolds.

The fact that the *RVE4/8* ortholog existed before the monocot-eudicot split also suggested that the *sMYB-PRR* linkage may represent an ancestral state. Scanning the *Amborella* genome revealed that ATR0661G529_ATR0661G570 (*PRR5/9*) and ATR0661G164 (*LHY/CCA1*) were co-located 18 genes apart, separated by 800 Kb on scaffold1 (Figure 2C). Likewise, ATR0594G076 (*PRR3/7*) and ATR0594G193 (*RVE4/8*) were co-located on scaffold23 separated by 27 genes and 900 kb (Figure 2C). Consistent with the organization in *Arabidopsis*, the linkages between the two *sMYB-PRR* clusters is present in *Amborella* suggesting that this linkage is an ancient arrangement of these core clock genes. Genetic linkages between *PHYA-PIF3*, *PHYB-LKP2* and *SRR1-BOA* were also identified in *Arabidopsis*, yet only *PHYA-PIF3* was also found in *Amborella* (Supplementary note; Supplemental Figure S3). Therefore, some of the these genetic linkages may result from associations forming only in *Arabidopsis*, while the fact that the *CCA1-PRR9*, *RVE4-PRR7* and *PHYA-PIF3* linkages date back to the basal plants suggests they could be evolutionarily significant.

### *sMYB-PRR* linkage inherited and closer in higher plants

Grape (*Vitis vinifera*) has been used extensively to unravel evolutionary relationships in the eudicot lineage since it only contains the ◻ WGD event (Jiao et al., 2014). If the *sMYB-PRR* linkages were evolutionarily significant then it might be expected that they are shared after distinct rounds of WGD. Syntenic orthologs between *Amborella* and grape were identified for both *CCA1/LHY-PRR5/9* and *RVE4/8-PRR3/7* pairs, with grape having 9 and 10 intervening genes respectively, fewer compared to the 18 and 27 respectively in *Amborella* (Figure 3A; Supplemental Figure S4). Whereas the *CCA1/LHY* and *RVE4/8* linkages are fractionated to one copy each, similar to *Arabidopsis*, *PRR5/9* and *PRR3/7* are retained in 3 and 2 copies respectively in grape (Supplemental Figure S4). These results suggest that there is a selective pressure to retain the *sMYB-PRR* linkage, possibly as a single copy each, and bring the linkage closer together.

**Figure 3.**
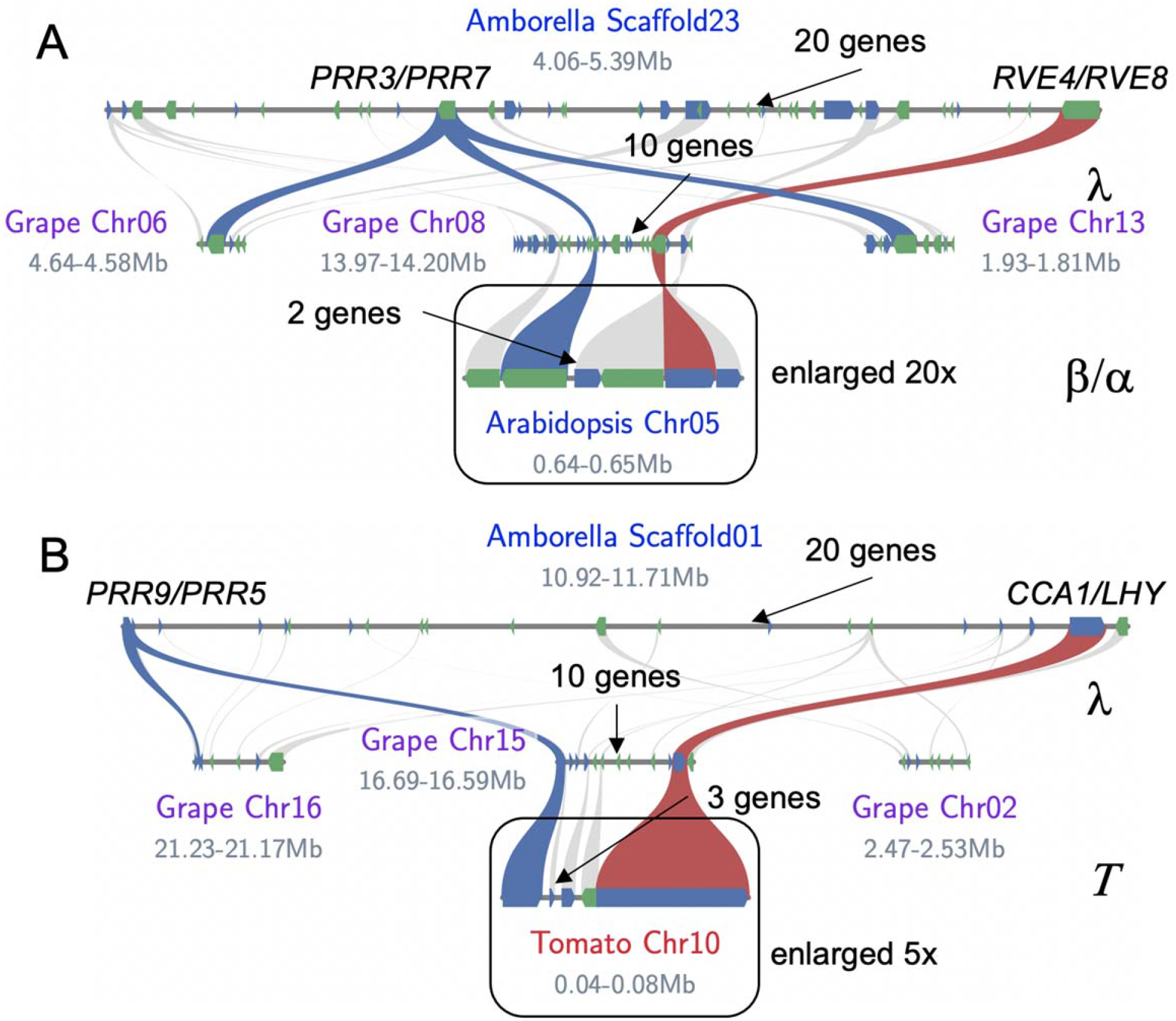
Syntenic sMYB-PRR pairs converge over evolutionary time. A) Syntenic blocks between *Amborella* (*Amborella trichopoda*), grape (*Vitis vinifera*) and *Arabidopsis* (*Arabidopsis thaliana*). There are three syntenic blocks between *Amborella* and grape as a result of the ◻ triplication. All three copies of *PRR3/7* (blue) are retained on grape Chr06, Chr08 and Chr13, while only one *RVE4/8* (red) is retained on Chr08, which results in the *sMYB-PRR* linkage 10 genes apart. Grape Chr08 is syntenic to *Arabidopsis* Chr05 where *RVE4-PRR7* are 2 genes apart; the *Arabidopsis* region is enlarged 20x to see the two intervening genes. B) Syntenic blocks between *Amborella* (*Amborella trichopoda*), grape (*Vitis vinifera*) and tomato (*Solanum lycopersicum*). Grape Chr08 is syntenic to tomato Chr10 where *RVE4-PRR7* are 3 genes apart; the tomato region is enlarged 5x to see the three intervening genes. The other syntenic genes (forward, blue; reverse, green) are depicted in the homologous chromosomal regions (grey).

*Arabidopsis* has experienced two additional WGD after the ◻ event shared with grape, which provides an additional opportunity to test how the *sMYB-PRR* linkage is evolving. Grape and *Arabidpsis* share one synthetic block between *RVE4/8* and *PRR3/7* (Figure 3A). The *CCA1/LHY* and *PRR5/9 linkage* does not exist between grape and *Arabidopsis* because grape does not have a *CCA1* ortholog, and the *LHY-PRR* association has been lost (fractionated) in *Arabidopsis* (Supplemental Figure 5). The *RVE4/8*-*PRR3/7* linkage is yet again reduced from 10 genes in grape to 2 genes in *Arabidopsis*. *Arabidopsis* is in the rosid clade of the angiosperms, as is grape, which could mean that the *sMYB-PRR* linkage is specific to this clade. Tomato (*Solanum lycopersicum*), which is in the asterid clade of angiosperms and has an independent WGD event (◻), was also found to retain the *sMYB-PRR* linkage with the distance between the *RVE4/8* and *PRR3/7* reduced to 3 genes (Figure 3B). Thus, not only is the *sMYB*-*PRR* linkage inherited from the basal *Amborella* lineage in syntenic blocks, but the genetic linkage also moves progressively closer together suggesting that during fractionation the *sMYB*-*PRR* linkage is preferentially retained.

### sMYB-PRR linkage is conserved across angiosperms except the Poaceae

Several plant genome databases pre-compute syntenic block information, which provides a broader view of whether the *sMYB-PRR* linkage is generally retained. The PLAZA 4.5 dicot database was searched for *sMYB* and *PRR* orthologous genes and the presence of linkages (Supplemental Table S5) (Van Bel et al., 2018). First, this analysis confirmed that only the Brassicacae have *CCA1*, while all other lineages lack *CCA1* and only contain *LHY* (Supplemental Figure S5). This result is consistent with the lack of *CCA1* in the Grape lineage and the fact that it arose in the progenitor to *Arabidopsis* as a result of the ◻ WGD event. Of the dicots, monocots, bryophytes, lycophytes and algae in the PLAZA 4.5 dicot database, 62% had at least one *sMYB-PRR* linkage. This value (62%) is probably a conservative estimate due to the quality of genomes and the possibility that the *sMYB*-*PRR* linkage could be more than 10 genes in some species; for example, the *Amborella* linkage is not detected in this analysis. 50% of the species had both *sMYB*-*PRR* linkages, while slightly more than half (56%) had only *RVE4/8-PRR3/7* linkages and only 19% of those had more than one of either *sMYB*-*PRR* linkage (Supplemental Table S5). A similar analysis with the PLAZA 4.5 monocot database revealed that only *Spirodela*, pineapple, palm and orchid retained the *sMYB-PRR* linkage while revealing that all grasses (Poaceae) tested have lost the linkage sometime after the sigma (◻) WGD shared by pineapple and the grasses (Figure 4; Supplemental Figure 6) (Ming et al., 2015; VanBuren et al., 2015). A recently published study leveraged a syntenic ortholog network approach across 123 high quality plant genome assemblies (Zhao et al., 2021), which when analyzed for the *CCA1LHY-PRR5/9*, *RVE4/8-PRR3/7* and *PHYA-PIF3* pairs provided additional evidence that these linkages were independently and convergently retained across the angiosperm lineage except the grasses (Supplementary note; Supplemental Table S6).

**Figure 4.**
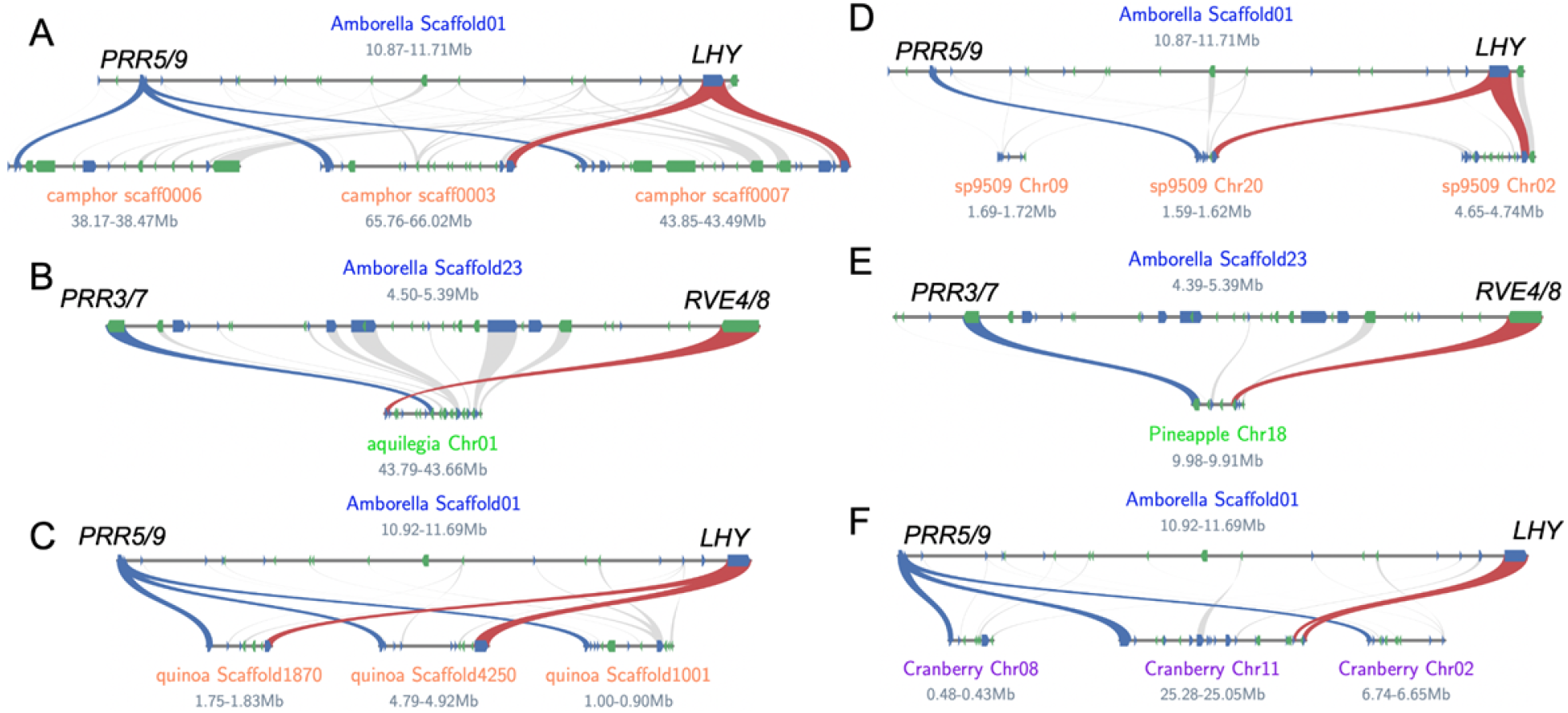
The *sMYB-PRR* linkage is conserved across Angiosperms. A) Two *LHY* (red) and *PRR5/9* (blue) linkages were retained in camphor (*Cinnamomum camphora*; Ranunculales). B) One *RVE4/8* (red)-*PRR3/7* (blue) was retained in aquilegia (*Aquilegia coerulea*; Magnoliids). C) Two *LHY* (red) and *PRR5/9* (blue) linkages were retained in quinoa (*Chenopodium quinoa*; Caryophyllales). D) One *LHY* (red)-*PRR5/9* (blue) linkage was found in sp9509 (*Spirodela polyrhiza* clone 5909; Araceae). E) One *RVE4/8* (red)-*PRR3/7* (blue) was found in pineapple (*Ananas comosus*; Commelinids). F) One *LHY* (red)-*PRR5/9* was found in cranberry (*Vaccinium macrocarpon*; Asterid); *LHY* is tandemly duplicated (TD) on Chr11. The other syntenic genes (forward, blue; reverse, green) are depicted in the homologous chromosomal regions (grey).

The syntenic ortholog network analysis revealed that many of the linkages across species were syntenic with *Amborella* despite distinct and numerous WGD. In general, the syntenic blocks detected between *Amborella* and other species are very small (just containing the *sMYB-PRR* linkage) suggesting that there is selective pressure to retain these genes in syntenic blocks even as surrounding genes are fractionated. For the *CCA1LHY-PRR5/9* and *RVE4/8-PRR3/7* linkages there are eight different combinations possible (Supplemental Figure 7), all of which were detected across an array of evolutionarily distinct monocots, dicots and basal plants: *Cinnamomum camphora* (Ranunculales), *Aquilegia coerulea* (Magnoliids), *Manihot esculenta* (Malpighiales), *Cuscuta australis* (Convolvulaceae), *Chenopodium quinoa* (Caryophyllales), *Vaccinium macrocarpon* (Asterid), *Apostasia shenzhenica* (Asparagales), *Ananas comosus* (Poales), *Cocos nucifera* (Arecales), *Elaeis guineensis* (Arecales) and *Spirodela polyrhiza* (Araceae) (Figure 4; Supplemental Figure 6; Supplemental Figure 7) (Zhang et al., 2017; Sun et al., 2018a; Ong et al., 2020; Kawash et al., 2021; Mansfeld et al., 2021; Yang et al., 2021b). In *Kalanchoe fedtschenkoi* (Saxifragales), a Crassulacean acid metabolism (CAM) photosynthesis plant that partitions carbon capture by TOD, the *LHY-PRR5/9* linkage has completely converged so the two genes are next to one another (no genes in between) (Yang et al., 2017). These results are consistent with the *sMYB-PRR* linkage being a general feature of flowering plant genome evolution.

### Serial retention in soybean of the *sMYB-PRR* linkage

In the broad analysis of *sMYB-PRR* linkages in the PLAZA 4.5 database, soybean (*Glycine max*) had the most linkages with a total of six: four *CCA1/LHY-PRR5/9* and two *RVE4/8-PRR3/7* (Supplemental Table S5); it has also retained all four *PIF3*-*PHYA* linkages (Supplemental Figure S3). Soybean has experienced two WGDs 59 mya and a recent one 13 mya; after these WGD events, 50% of genes are still retained in syntenic pairs (Zhao et al., 2017). However, soybean has retained more than triple of the sMYB-PRR linkages compared to its close relatives *Cicer arietinum* (chickpea), *Medicago truncatula* (barrel clover) and *Vigna radiata* (mung bean), which have retained one of each for a total of two (Supplemental Table S5).

Like other species outside of the Brassicaceae, soybean does not have *CCA1* or *RVE4*, yet has four copies of *LHY* and *RVE8*, in addition to two copies of *PRR7* and 4 of *PRR9*. Syntenic block analysis revealed that these genes formed the six different *sMYB-PRR* linkages (Figure 5). Leveraging Ks to date the syntenic blocks revealed that both linkages were retained after the WGD 59 mya, and that the *LHY-PRR9* linkage was retained after the WGD 13 mya while the *RVE8-PRR7* was fractionated. The similarity in fractionation between Chr03 and Chr19 suggests that the *PRR7* was fractionated before the WGD 13 mya that resulted in the retention of *RVE8* but the loss of two *sMYB-PRR* linkages. While soybean retains all four syntenic linkages of *PIF3*-*PHYA* the blocks have significantly diverged since the WGD 59 mya resulting in the linkages on Chr10 and Chr20 being 7 and 10 genes apart respectively and the linkages in Chr03 and Chr19 being four genes apart (Supplemental Figure S3). *PHYA* has been reported to be the gene underlying two soybean maturity group (MG) loci E3 and E4 (Nissan et al., 2021), which were found on Chr19 and Chr20 and separated by 56 mya.

**Figure 5.**
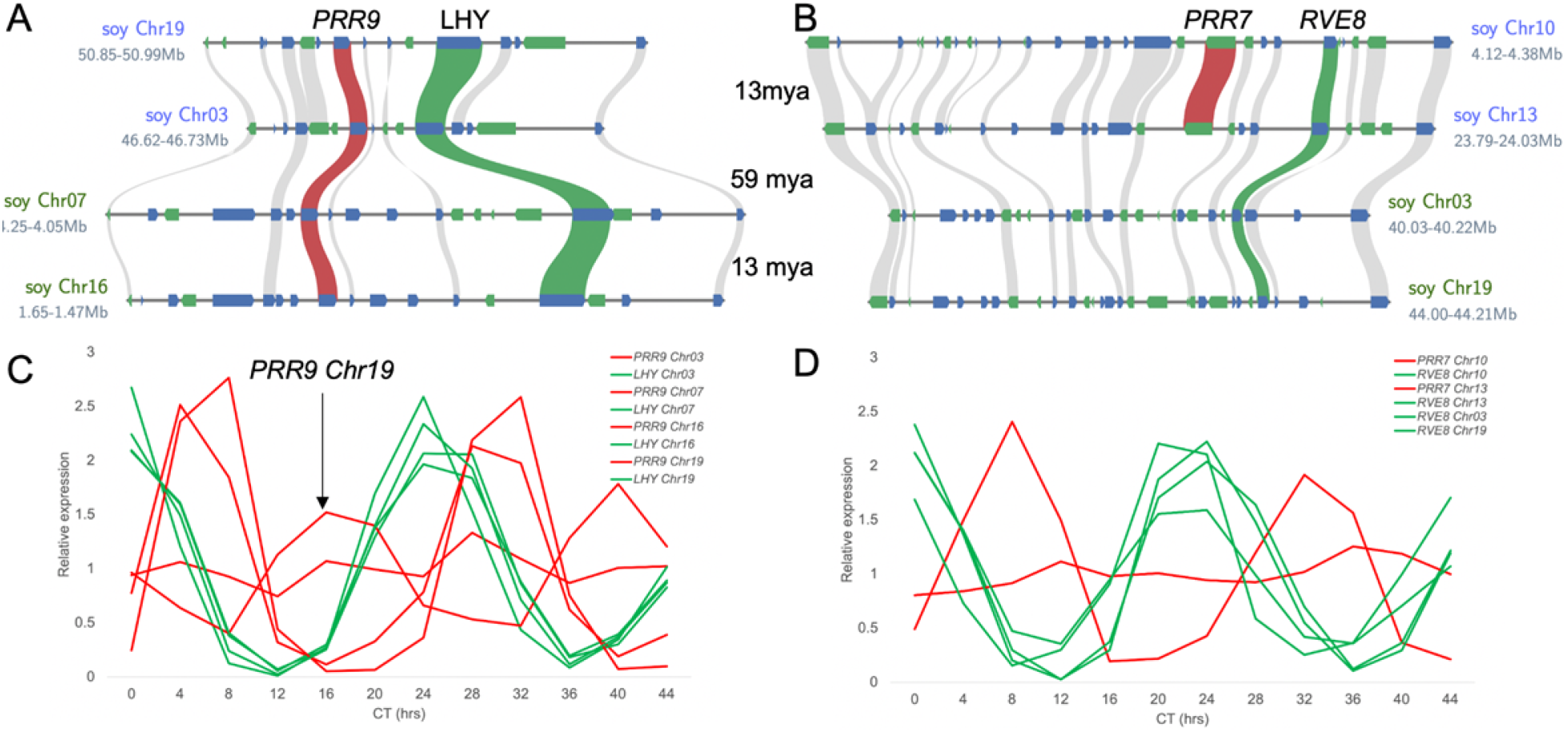
The *sMYB-PRR* linkages are preferentially retained in soybean. A) In Soybean (*Glycine max*; Williams82) the *LHY* (green) and *PRR9* (red) linkage is retained over two whole genome duplications (WGDs) 59 million years ago (mya) between Chr03-Chr07 and 13 mya between Chr03-Chr19 and Chr07-Chr16. B) In contrast, only two *RVE8* (green) and *PRR7* (red) linkages are retained on Chr13-Chr10 after the most recent WGD 13 mya, while the other two copies are lost due to fractionation. The other syntenic genes (forward, blue; reverse, green) are depicted in the homologous chromosomal regions (grey). C) Expression of the soy *PRR9* (red) and *LHY* (green) and D) *PRR7* (red) and *RVE8* (green) orthologs in continuous light and temperature over 44 hrs (Circadian Time; CT).

Reanalysis of a recently published RNA-seq circadian time course in soybean revealed that while all of the *sMYB* (*LHY* and *RVE8*) in the linkages retained the morning-specific phase (CT0) similar to *Arabidopsis*, the phases of the *PRRs* were variable (Figure 5C, D). Specifically, the *PRR9* on Chr16 is expressed in the evening (CT17) compared to the morning expression of the other paralogs (CT6) (Figure 5C). Only the *PIF3*-*PHYA* linkage on chr20, which is the MG E4 locus, cycles with a similar TOD expression as *Arabidopsis* with *PHYA* peaking at dusk (CT8) and *PIF3* peaking in the middle of the night (CT17) (Supplemental Figure S8). The other *PIF3*-*PHYA* linkages, including the MG E3 locus on Chr19, have very low expression and are not predicted to cycle. In contrast to the *LHY/CCA1*-*PRR5/9* and *RVE4/8-PRR3/7* linkages that have morning to midday expressions, the *PIF3*-*PHYA* linkage peaks midday to midnight. Recently it was shown that knocking out all four of the *LHY* paralogs using CRISPR-CAS9 impacts plant architecture resulting in smaller plants and reduced internode length (Cheng et al., 2019). These results coupled to the preferential retention of the sMYB-PRR linkage over two rounds of WGD suggest this association could be of importance and a target for future soybean development via chronoculture (Steed et al., 2021).

### Maize and the loss of *sMYB-PRR* linkage

The loss of the *sMYB-PRR* linkage in the economically and agriculturally important grasses provides an opportunity to probe the significance of the association. *Oropetium* has one copy each of *LHY*, *PRR5/9* and *RVE8*, while it retains two copies of *PRR7*, all on different chromosomes (Figure 6; Supplemental Figure S6). The same is true for rice and *Sorghum*, except that *Sorghum* only has one *PRR7* copy (Figure 6A); similar results were found for *Setaria*, *Brachypodium*, and other grasses for which high quality genomes exist. However, maize (*Zea mays*; B73), which has experienced a recent WGD, retains two copies of each gene and has a slightly different pattern with only one *RVE8* (Figure 6A). A closer look compared to *Oropetium* revealed that not only was the second *RVE8* fractionated but *PRR7* matched the two *Oropetium* chromosomal locations and the second copy of *PRR7* was fractionated (Figure 6B,C). Since maize lines have a high level of presence absence variation (PAVs) (Springer et al., 2009; Sun et al., 2018b), it is possible that the fractionation of these genes represents real differences in the content of *sMYB* and *PRR* between lines.

**Figure 6.**
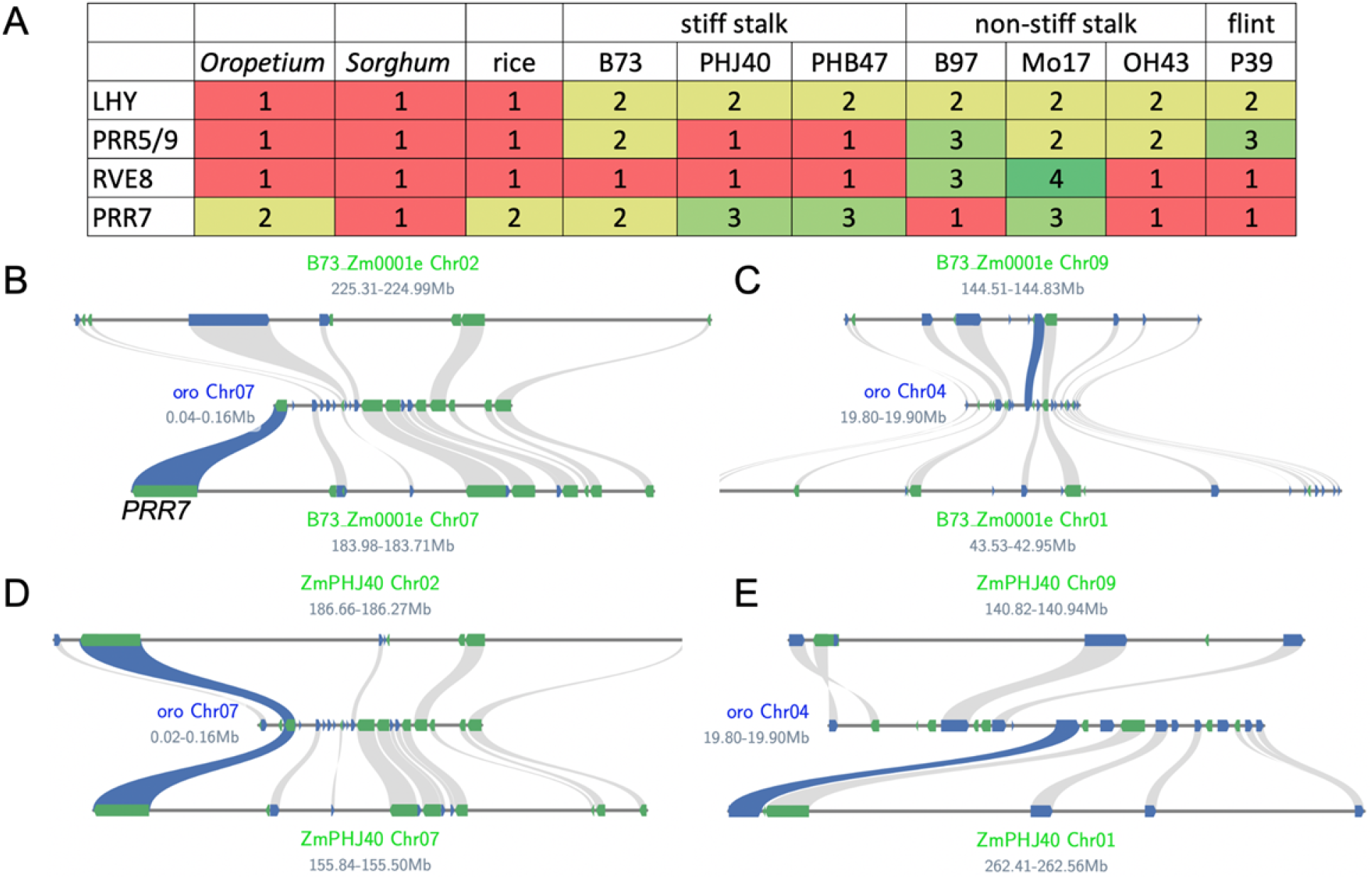
Presence absence variation (PAV) of the *sMYB-PRR* genes in maize heterotic groups. A) Number of *LHY*, *PRR5/9*, *RVE8* and *PRR7* copies across *Oropetium* (oro; *Oropetium thomaeum*, *Sorghum* (*Sorghum bicolor*), rice (*Oryza sativa*), stiff stalk maize (B73, PHJ40 and PHB47), non-stiff stalk maize (B97, Mo17, OH43) and flint maize (P39). Squares are colored to draw contrast to the numbers: >1 (green) and =1 (red). B-C) The four maize (B73; Zm0001e) syntenic regions with the two *PRR7* (blue) copies found in *Oropetium* (oro) on Chr07 (B) and Chr04 (C). B73 only has two copies of *PRR7* on Chr07 and Chr09; the other two have been lost through fractionation. D-E) The maize line PHJ40 (ZmPHJ40) has 3 copies of *PRR7*; two copies are retained in syntenic blocks with oro on Chr07 and Chr02, the latter of which has been lost in B73 (B). One *PRR* copy is found in the oro Chr04 syntenic block on Chr01, which is the opposite found in B73 (C), consistent with all four copies of *PRR7* segregating in heterotic groups.

Maize lines have been developed into specific inbred heterotic groups such as stiff stalk (SS), non-stiff stalk (NSS) and flint (F) that are marked by high levels of PAVs, and when crossed, form the commercial hybrids that display heterosis resulting in high yields and large plants (Bornowski et al., 2021). Looking at the *sMYB* and *PRR* genes across each of the heterotic groups in high quality maize genomes revealed PAVs in these clock genes except *LHY*, which always had two copies resulting from the most recent WGD (Figure 6A) (Hirsch et al., 2016; Sun et al., 2018b; Bornowski et al., 2021; Hufford et al., 2021). While the majority of maize lines retain two copies of *PRR7* syntenic to *Oropetium*, some retain three as a result of a retention on Chr02; PHJ40 is unique in that it retains the copy on Chr01 instead, which suggests that all four *PRR7* copies resulting from the most recent WGD may be differentially segregating across maize heterotic groups. In contrast, *RVE8* has one syntenic copy across all lines tested, but the additional copies in the NSS (MO17 and B97) are interspersed, suggesting they were duplicated in a manner other than WGD. When there are two copies of *PRR5/9,* they are syntenic pairs resulting from the most recent WGD, while only one copy represents a fractionation (PHJ40 and PHB47) and three copies (B97 and P39) is the result of a non-syntenic dispersed duplication. These results suggest the ability in maize to inherit different versions of the *sMYB/PRR* paralogs is important in grasses and provides a clue as to why the *sMYB-PRR* linkage was broken.

### Origin of the *sMYB-PRR* linkage

Recently it was shown that *RVE8* and *LHY* genes date as far back as unicellular green algae, and that these genes have antagonistic roles in the clock’s response to the environment (Shalit-Kaneh et al., 2018). The circadian clock in *Ostreococcus tauri*, a green unicellular picoalage, is controlled by a simple two component negative feedback loop of one *sMYB* and *PRR* (Corellou et al., 2009). In *O. tauri* and the closely related *O. lucimarinus*, there is one sMYB that clusters with *LHY* and two that cluster with *RVE* (Supplemental Figure S9). *O. tauri* has one *PRR* gene, while *O. lucimarinus* has two copies, which were the result of a WGD, all of which have been called *PRR1-like* (Corellou et al., 2009). However, based on the gene trees, they are situated between the *PRR1* and *PRR5/9* clades (Supplemental Figure S9). All five genes are located on different chromosomes in both *O. lucimarinus* and *O. tauri*, indicating that even though both the *sMYB* and *PRR* clades are present, these two algae do not share the *sMYB*-*PRR* linkage (Figure 7). This is also true in the other chromosome resolved algae such as *Micromonas pusilla* (CCMP1545) and *Chlamydomonas reinhardtii*, both of which have two *sMYB* and one *PRR*, but they are found on separate chromosomes (Figure 7).

**Figure 7.**
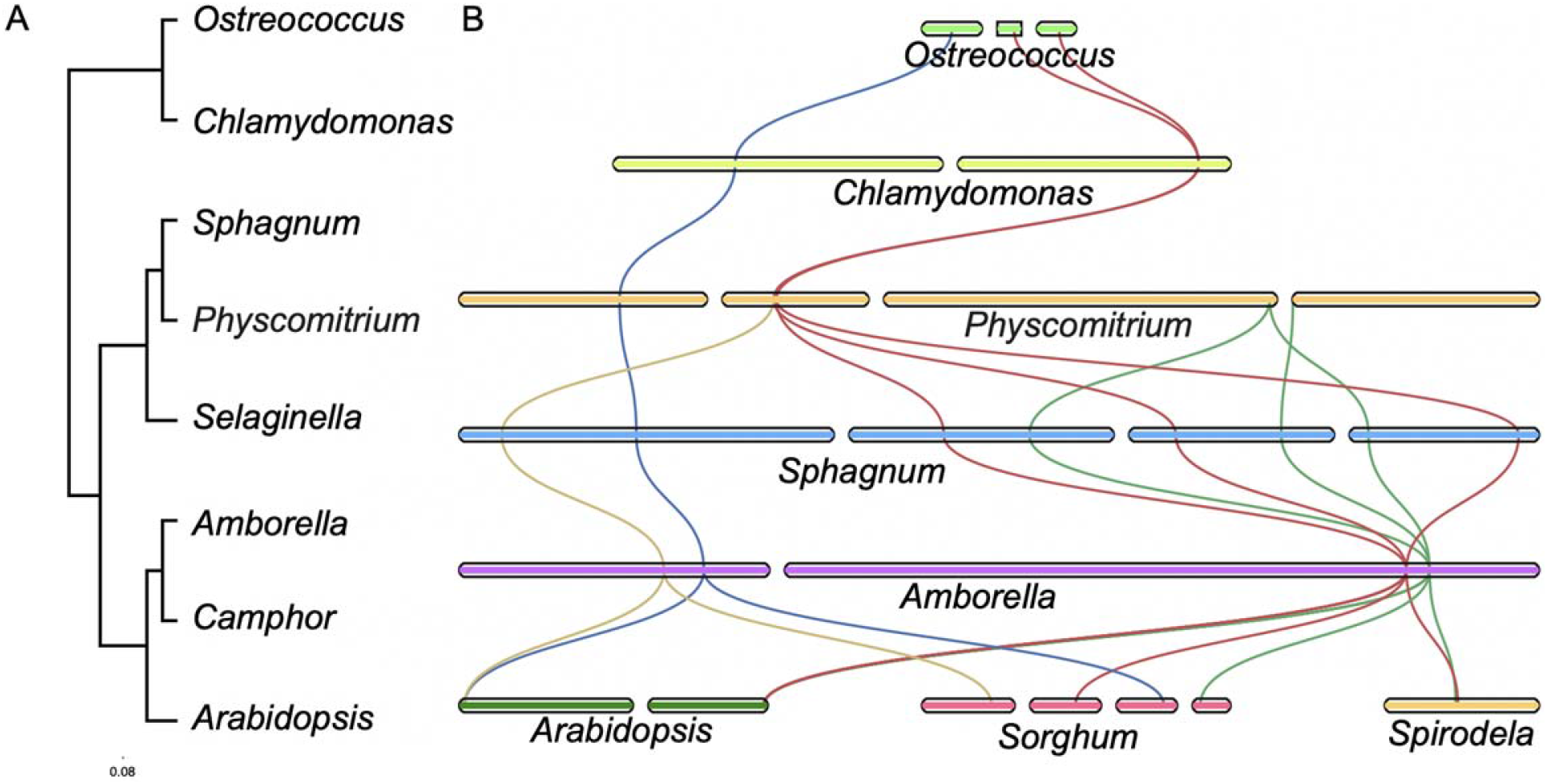
The *sMYB-PRR* linkage arose in the bryophyta. A) Phylogenetic tree spanning the green tree of life from green algae to higher plants. Green algae: *Ostreococcus* (*Ostreococcus lucimarinus*), and *Chlamydomonas* (*Chlamydomonas reinhardtii*); Bryophytes and Lycophytes: *Sphagnum* (*Sphagnum angustifolium*), *Physcomitrium* (*Physcomitrium patens*), and *Selaginella* (*Selaginella lepidophylla*); and higher plants *Amborella* (*Amborella trichopoda*), *Camphor* (*Cinnamomum camphora*), and *Arabidopsis* (*Arabidopsis thaliana*) B) chromosomal view of the *RVE4/8* (blue), *CCA1/LHY* (green), *PRR5/9* (red) and *PRR3/7* (yellow) over evolutionary time. *Sorghum* (*Sorghum bicolor*) and *Spirodela* (*Spirodela polyrhiza*).

The charophyte (green algae), bryophyte (liverwort, hortwort, moss), and lycophtes (spike-moss) lineages all include both *sMYB* clades and *PRRs (Linde et al., 2017; Li et al., 2018; Ferrari et al., 2019; Wickell et al., 2021)*. However, no *sMYB*-*PRR* linkage was detected in these genomes at either short or long distance; many of these genome assemblies are not chromosome-resolved, which could mean the linkage would be missed if it were on the scale found in *Amborella*. In addition, two chromosome-resolved moss genomes of *Physcomitrium patens* (formerly *Physcomatrella patens*) and *Ceratodon purpureus* have the *sMYB* and *PRR* on separate chromosomes and therefore clearly unlinked (Lang et al., 2018; Carey et al., 2021). A chromosome-resolved genome assembly of a bryophyte is available for *Sphagnum angustifolium (formally Sphagnum fallax*), which provides an opportunity to evaluate a third moss genome (Meleshko et al., 2021). There was an expansion of both the *sMYB* and *PRR* gene families with seven and five respectively, four of which are located on the same chromosome (Figure 7). There are three *LHY/CCA1*-*PRR37* linkages on linkage group (LG)04, LG10, and LG15, that are 8, 8 and 12 Mb apart respectively. The fourth is a *RVE6*-*PRR37* combination 12 Mb apart on LG02 (Figure 7). A similar situation was observed for *PIF3-PHYA*, where three pairs were found 8, 7, and 7 Mb apart on LG06, LG07 and LG12. While these are not closely linked like in *Arabidopsis*, the fact that they are on the same chromosome much like *Amborella* is suggestive that this may represent a linkage that predated the embryophyta (land plants).

Only recently have fern genomes (Li et al., 2018) and high-quality gymnosperm genomes (Scott et al., 2020) become available to assess the core circadian clock genes in the context of a genome. The *Azolla* and *Salvinia* genomes contain both *CCA1/LHY* and *RVE* clades with one and three respectively, as well as *PRR1* and *PRR3/7*. However, none of these are in linkage on the contigs/scaffolds; once again this is possibly due to the fact they are not assembled at a chromosome scale and they may be at the same or greater distance (~12 Mb) as *Sphagnum* (*S. angustifolium*). A similar problem is encountered with gymnosperm genomes since they are between 5-30 Gb in size (Michael, 2014). However, it has been shown that the core circadian genes are conserved in conifers (Nose and Watanabe, 2014). Recently, a high-quality chromosome-resolved genome 8 Gb in size has become available for the Giant Sequoia (*Sequoiadendron giganteum*) (Scott et al., 2020). The Sequoia genome has a *RVE4/8*-*PRR3/7* linkage 12 Mb apart (51 genes) on Chr07 similar to that found in *Sphagnum* and collinear with *Amborella* (Supplemental Figure S7). Along with *Sphagnum*, the presence of *sMYB* and *PRR* on the same chromosome in the gymnosperm suggests that the linkage emerged before flowering plants.

Taken together these results show that the early algae had both clades (*CCA1*/*LHY* and the *RVEs*) of the *sMYBs* but the *sMYBs* and *PRRs* were found on separate chromosomes. In some bryophytes and the gymnosperms *sMYB* and *PRR* genes were found on similar chromosomes yet at a distance (~12 mb), suggestive of an emerging *sMYB*-*PRR* linkage. In the basal angiosperm *Amborella* the *sMYB*-*PRR* linkage is closer (~1 mb) and is inherited across the entire flowering plant lineage, except the grasses (Poaceae) (Figure 8).

**Figure 8.**
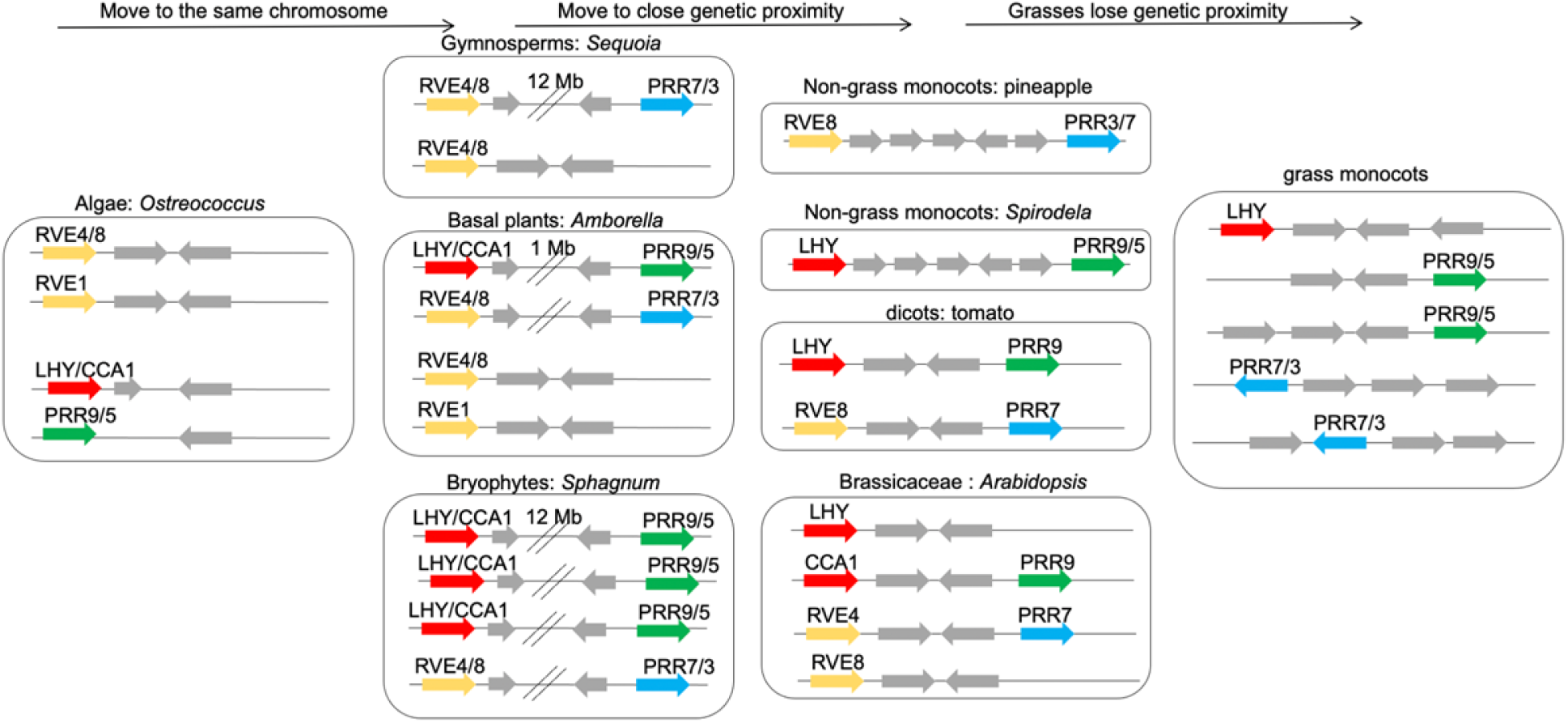
Model of the *sMYB-PRR* linkages across the green lineage. The Green alga (*Ostreococcus lumimarinus*) has *RVE4/8* and *RVE1* (yellow), *LHY/CCA1* (red) and *PRR9/5* (green), and these are all found on separate chromosomes; intervening genes are included (grey). Starting with *Sphagnum* (*Sphagnum angustifolium*) in the bryophyte lineage *sMYB* and *PRR* gene combinations are found on the same chromosomes 12 megabases (Mb) apart.

Similarly, the *RVE4/8*-*PRR3/7* linkage is found with Giant Sequoia (*Sequoiadendron giganteum*) also 12 Mb apart on the same chromosome. In the basal angiosperm *Amborella*, both *sMYB*-*PRR* linkages are found closer together at 1 Mb. In higher plants such as *Arabidopsis* (*Arabidopsis thaliana*) both *sMYB*-*PRR* linkages are found 2-3 genes apart; the Brassicaceae is the only lineage with *CCA1*, the result of a WGD, and the *LHY-PRR5/9* linkage is lost. Non-grass monocots, such as pineapple (*Ananas comosus*) and Spirodela (*Spirodela polyrhiza*) retain one of the *sMYB*-*PRR* linkages with a variable number of intervening genes, while most major angiosperm clades retain both *sMYB*-*PRR* linkages (Supplemental Table S6 and Supplemental Table S7). One exception are the grasses (Poaceae) that have lost the *sMYB*-*PRR* linkages and the four genes are found on separate chromosomes.

## Discussion

Here it is shown that the *LHY*/*CCA1-PRR5/9*, *RVE4/8-PRR3/7* and *PIF3-PHYA* genetic linkages reflect ancient associations that emerged as early as the bryophyte lineage. *CCA1* emerged from *LHY* after the lambda triplication in dicots, and was also lost (fractionated) in all lineages except the Brassicacae, which explains why it is not found in monocots or most higher plants. The basal angiosperm *Amborella* genome has all three genetic linkages at about 1 Mb apart, which are brought in tighter linkage across all lineages of higher plants despite distinct WGD and fractionation events. Some genetic linkages are very close such as *Arabidopsis* and the CAM plant *Kalanchoe* that have two and zero intervening genes respectively. While the all three linkages are maintained in the non-grass monocots, the grasses have lost the linkages, and *sMYB*, *PRR*, *PIF3* and *PHYA* genes are found on different chromosomes. In contrast, soybean has preferentially retained 10 of 12 linkages, only losing two *RVE8-PRR7* linkages after two rounds of WGD and fractionation, suggesting that the linkage is important in a modern day crop species.

The genetic linkage of the core circadian and light signalling genes suggests there is selective pressure to keep these components close genetically. In general, genes that are proximal in chromosome space (genetically linked) segregate together due to the low probability of a crossover event occurring between them; the closer genes are the more likely they will segregate together. To arrive at this state there are several factors in play. First, higher plant genomes are highly variable due to their ability to hybridize (polyploidy), duplicate and reduce (fractionate) to a diploid state (Qiao et al., 2019). The *sMYB*-*PRR* and *PIF3-PHYA* linkages were retained across all major clades angiosperms despite variable intervening gene numbers, ranging from from 51 genes in Sequoia to ~18 genes in *Amborella*, ~9 genes in grape, 2 in Arabidopsis, and none in *Kalanchoe*. *Kalanchoe* is an obligate CAM photosynthesis plant, which means that carbon capture is regulated in time-of-day (TOD) fashion, suggesting that the intimate *LHY-PRR5* linkage is of functional significance.

In addition to the genetic linkages moving closer together over distinct WGD events and lineages, only a limited number of specific linkages are retained. Most (>70%) lineages retained one copy of each *sMYB*-*PRR* linkage or the *PIF3-PHYA* linkage, and only 19% had more than one copy of any of the linkages (Supplemental Table S5 and Supplemental Table S6). This result suggests that having one copy of each linkage is the preferred state, and could reflect the “dosage hypothesis” that there is a pressure to balance expression among genes that are highly networked or which encode members of multi-protein complexes (Birchler and Veitia, 2012; Lou et al., 2012). This is also seen in other core clock genes such as *GIGANTEA* (*GI*) and *TIMING OF CAB EXPRESSION1* (*TOC1/PRR1*) that are found as single copy genes across most lineages, consistent with core clock components being fractionated to a limited copy number to retain proper dosage. Exceptions exist for components of the clock in crop species that retain more copies of clock genes, such as soybean that retained 10 of 12 linkages over its last two WGD events (discussed further below).

The results presented here suggest that *LHY* and *RVE8* represent the core ancestral *sMYB* clock components, which is consistent with a recent study in *Arabidopsis* that provides some clues as to the significance of the two *sMYB-PRR* linkages (Shalit-Kaneh et al., 2018). While the triple *rve4/6/8* and double *cca1/lhy* mutants result in long and short periods under continuous conditions respectively, the quintuple mutant has a near wild type period and phenotypes (Shalit-Kaneh et al., 2018). These results are consistent with *CCA/LHY* acting as repressors and *RVE4/6/8* acting as activators (Figure 1A) (Farinas and Mas, 2011; Rawat et al., 2011; Hsu et al., 2013). In addition, it shows that these individual feedback loops are not necessary for the generation of circadian rhythms. However, the quintuple mutant displays a loss in robustness and temperature compensation suggesting that these loops are critical for clock sensitivity to environmental input. Therefore, it is possible that the tight linkage between the *sMYB* and *PRRs* ensures a specific balance of negative and positive feedback loops to maintain robust responses to daily fluctuating environmental variables.

By extension, this hypothesis of the significance of the *sMYB*-*PRR* linkage would suggest soybean has a specific robustness to environmental variables. The initial retention after WGD (~59 mya) of the first four linkages most likely pre-dated human selection in the soybean native range of Asia, while the most recent (~5-13 mya) WGD resulting in retention of ten of the potential twelve combinations could have been the result of breeding. Soybean is particularly interesting in terms of its domestication and the circadian clock because it has been bred into maturity groups (MGs) to enable it to grow under an array of latitudes and environmental conditions(Li and Lam, 2020). There are at least 12 loci underlying MGs, of which two are core circadian clock genes (*EARLY FLOWERING 3*; *ELF3a* and *GIa*), three are flowering time genes (*FLOWERING LOCUS T*; FT3a; FT1a and TERMINAL FLOWER1; TFL1b) and two are the *PHYA* genes discussed (*PHYA3, PHYA3*),(Nissan et al., 2021). The results presented here suggest that the E4 MG might be an evening specific association between *PHYA* and *PIF3*, and not just PHYA, which could provide a balance of alleles since it is known that PIFs play a role in negatively regulating the phytochromes. In addition, it has been shown through quantitative trait loci (QTL) and genome wide association studies (GWAS) that *PRR3* is a target of domestication as well (Li et al., 2019b; Lu et al., 2020).

The fact that the grasses have lost the linkages is confounding but also provides clues as to their significance. The commelids clearly have the *sMYB-PRR* association as seen in pineapple, and then it is lost sometime after the sigma (◻) WGD and fractionation of *Oropetium* (Figure 4; Supplemental Figure S6). Looking at *sMYB* and *PRR* orthologs in maize revealed that while *LHY* and *PRR5/9* are retained in duplicate after the recent WGD, there are four *PRR7* with different PAV across several heterotic groups (Figure 6). It has been shown that circadian clock plays a role in hybrid vigor, or heterosis in rice, maize and *Arabidopsis* (Ni et al., 2009; Shen et al., 2015; Ko et al., 2016; Yang et al., 2021a), and this PAV of *PRR7* could play a role in the hybrid vigor seen when heterotic groups are crossed. There is some precedence that PRR7 plays a role in agronomically important traits in the grasses; In rice, *PRR3/7* is *HEADING DATE 2* (*HD2*) and variation at this locus provides photoperiod and temperature sensitivity (Koo et al., 2013). Maize as well as other grasses show particularly strong hybrid vigor, which may result in the selection states where genes are unlinked to optimize for the exploration of more ecological niches.

## Conclusion

A recent report provides additional important clues as to the significance of the genetic linkages of the core circadian and light signaling genes. It was shown that the waves of *PIFs* and *PRRs* physically interact to control growth-related genes at dawn (Martín et al., 2018). With this paradigm one could imagine that the *LHY*/*CCA1-PRR5/9*, *RVE4/8-PRR3/7* and *PIF3-PHYA* genetic linkages are maintained to ensure a very specific feedback loop based on “defined” reciprocal effectors for a specific environment. Since plants cannot move from the environment they germinate in, it is critical that they have an innate sense of time.

## Material and Methods

### Circadian clock genes

A list of core circadian clock genes, as well as light signaling and flowering time genes associated with the clock were used to seed the original analysis (Supplemental Table S1) (Lou et al., 2012).

### Genomes

Genomes described were downloaded from Phytozome13 (https://phytozome-next.jgi.doe.gov/) and PLAZA (https://bioinformatics.psb.ugent.be/plaza/). When genomes were downloaded from other sources the primary reference was cited at the first mention of the species in the text.

### Gene family analysis

Proteins for species specifically mentioned in the text were evaluated for orthology using orthofinder (Emms and Kelly, 2015). Different combinations of species were run during multiple rounds of orthofinder to detect when the sMYB and PRR family relationships merged and split. In general, smaller clustering of closely related species resulted in the PRRs splitting into separate families (*PRR1* and the other PRRs), as well as *CCA1* (when it was present in the Brassicaceae)/*LHY* and the RVEs. Families were manually compared to the results in the PLAZA databases: PLAZA dicot 4.5 and monocot 4.5 databases (Van Bel et al., 2018).

### Syntenic ortholog analysis

All syntenic analyses were conducted with either MCscan using the python version (https://github.com/tanghaibao/jcvi/wiki/MCscan-(Python-version) or CoGe SynMap (Grover et al., 2017). All syntenic ortholog searches were conducted by all-by-all protein alignments using last. Both MCscan and DAGchaniner (CoGe) were run with 20 genes max blocks with 5 syntenic pair minimum. Syntenic plots were generated using MCscan python version. Broad syntenic relationships across plant genomes were analyzed using the PLAZA dicot 4.5 and monocot 4.5 databases (Van Bel et al., 2018), as well as syntenic relationships identified across 123 high quality plant genomes using a network approach (Supplementary note) (Zhao et al., 2021).

### Expression analysis

Published *Arabidopsis* (Yang et al., 2020) and soy (Li et al., 2019a) RNA-seq datasets were downloaded from the Sequencing Read Archive (SRA) and reanalyzed using the updated DIURNAL pipeline (https://gitlab.com/NolanHartwick/super_cycling) using the same reference genomes described in this study (Michael et al., 2020; Wickell et al., 2021). The resulting cycling matrix was used to plot cycling expression profiles.

## Acknowledgements

Thank you to the Michael Lab for helpful comments and discussion on the manuscript. Also, special thanks to C. Robertson McClung for inspiring this work and providing critical reading and feedback. Thank you to Jon Shaw, David Weston and Jeremy Schmutz for the use of the *Sphagnum angustifolium* (formally called *Sphagnum fallax*) genome prior to their genome publication.

## Supplementary note

### *PHYA-PIF3* linkage

In addition to the *CCA1-PRR9* and *RVE4-PRR7* linkages, other clock and light associated genes were searched for close linkages. From the initial list of 61 genes we found 5 linkages that were within 13 genes: *CCA1-PRR9* (4), *RVE4-PRR7* (3), *PHYA-PIF3* (4), *PHYB-LKP2* (12) and *SRR1-BOA* (1). Looking at both *Amborella* and soy as two distinct reference points in evolutionary history, these three additional linkages were tested for proximity. Only *PHYA-PIF3* linkages were retained in both *Amborella* and soy, and looking at the progression through several WGD, the *PHYA-PIF3* progressively gets closer as seen with *LHY/CCA1-PRR5/9* and *RVE4/8-PRR3/7* (Supplemental Figure S3). Similar to the *LHY/CCA1-PRR5/9* and *RVE4/8-PRR3/7* linkages, both *PHYA* and *PIF3* cycle with slightly different phases of ZT11 and ZT16 respectively (Supplemental Figure S9), which in contrast makes them distinctly evening-specific compared to *LHY/CCA1-PRR5/9* and *RVE4/8-PRR3/7*. The same is also true of one of the four *PHYA-PIF3* syntenic pairs found in soy; *PHYA* and *PIF3* have antiphasic expression at CT8 and CT16 (Supplemental Figure S9B).

### Microsynteny across 123 plant species

A comprehensive dataset was published recently that looked at the microsynteny of 123 plant species spanning angiosperms using a novel network approach, which provided an additional opportunity to validate and extend the clock gene linkages (Zhao et al., 2021). The goal of the network approach was to generate a binary array of “syntenic orthologs” for deep phylogenetic inference, which resulted in gene families that were restricted to only genes in syntenic blocks. Syntenic gene families were searched for the *Arabidopsis CCA1, LHY, RVE4, RVE8, PRR3, PRR5, PRR7, PRR9, PIF3* and *PHYA*. Consistent with the results presented with other methods, *CCA1-LHY*, *RVE4-RVE8*, *PRR3-PRR7*, and *PRR5-PRR9* were in syntenic families 3634, 4022, 1555 and 1018 respectively. *PIF3* and *PHYA* were also in syntenic families, and while the *Arabidopsis PHYA* is alone in syntenic family 6934 (this means it has been reduced to one copy in *Arabidopsis* through fractionation), *PIF3* is part of a family that includes *Arabidopsis HFR1*, *PIL1* and *PIL2* in syntenic family 851 (Supplemental Table S6 and Supplemental Table S7).

The *PRR3-PRR7* and *PIF3* families contained 98.4% and 96.7% of the 123 plant species in syntenic blocks respectively (Supplemental Table S7). This places these syntenic families in the top 0.5% of all families for the number of species represented, which means the blocks that contain these genes are highly conserved. In addition, both families share blocks with the basal angiosperm *Amborella*, which is only found in 7.6% of families, making the *PRR3-PRR7* and *PIF3* families not only conserved but also rooted in the basal angiosperm lineage. The other families (*CCA1-LHY*, *RVE4-RVE8*, *PRR5-PRR9* and *PHYA*) are also in the 7.6% of families rooted in *Amborella* (share syntenic blocks) consistent with the fact that all of the families are also highly conserved over time. However, in contrast these other families only had syntenic blocks in between 69% and 81% of the 123 plant species. The absence of syntenic blocks in these families compared to *PRR3-PRR7* and *PIF3* was primarily driven by the lack of these genes being found in syntenic blocks in the grasses (Poaceae) (Supplemental Table S6). The *PHYA* family also was not found in syntenic blocks in the other monocots as well as the Solanaceae, Cucurbitaceae, and Malvaceae. Most species had more than one syntenic block for all of the families consistent with a history of WGD and retention of these blocks. The median for *CCA1-LHY*, *RVE4-RVE8*, and *PHYA* was 1 syntenic block per species while *PRR3-PRR7*, *PRR5-PRR9* and *PIF3* was 2 syntenic blocks per species.

Although a high percentage of the species have the clock genes in syntenic blocks, the *LHY/CCA1-PRR5/9*, *RVE4/8-PRR3/7*, *PIF3-PHYA* linkages may have been lost due to differential fractionation, yet 61%, 56% and 57% retained at least one and 11%, 9% and 12% retained more than one linkage respectively (Supplemental Table S6). While many of the species with multiple linkages are polyploid (cotton, camelina, etc), soy has the highest number of retained linkages across all three gene sets as found in other datasets. 73% of species have at least one of the *LHY/CCA1-PRR5/9* and *RVE4/8-PRR3/7* linkages, and 53% have one of these linkages as well as the *PIF3-PHYA* linkage; 34% of species have all three linkages. Similar to the analysis from other datasets, the grasses (Poaceae) have completely lost the associations despite *PRR3/7* and *PIF3* being found in syntenic blocks. Considering that the grasses make up 13% of the species, the tight linkage between clock genes is found in a high number of non-grass species (Supplemental Table S6).

### Supplemental Figures

**Supplemental Figure S1.**
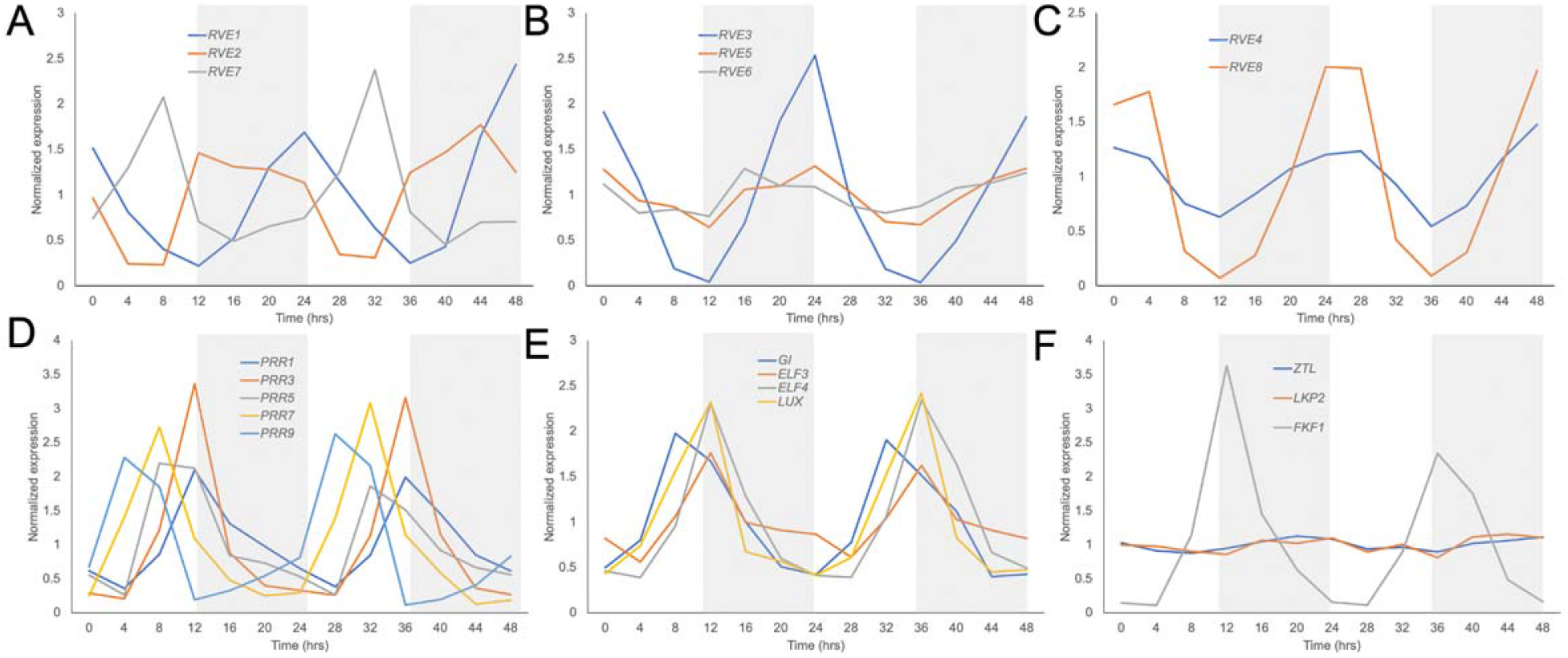
Expression of core circadian clock genes in *Arabidopsis*. Core circadian clock genes grouped by gene family or function. A) *RVE1*, *RVE2* and *RVE7* B) *RVE3*, *RVE5*, and *RVE7* C) *RVE4* and *RVE8*; D) *PRR1*, *PRR3*, *PRR5*, *PRR7* and *PRR9*; E) *GI*, *ELF3*, *ELF4*, and *LUX*; F) *ZTL*, *LKP2* and *FKF1*. Normalized RNA-seq expression was plotted over the day with grey boxes representing dark.

**Supplemental Figure S2.**
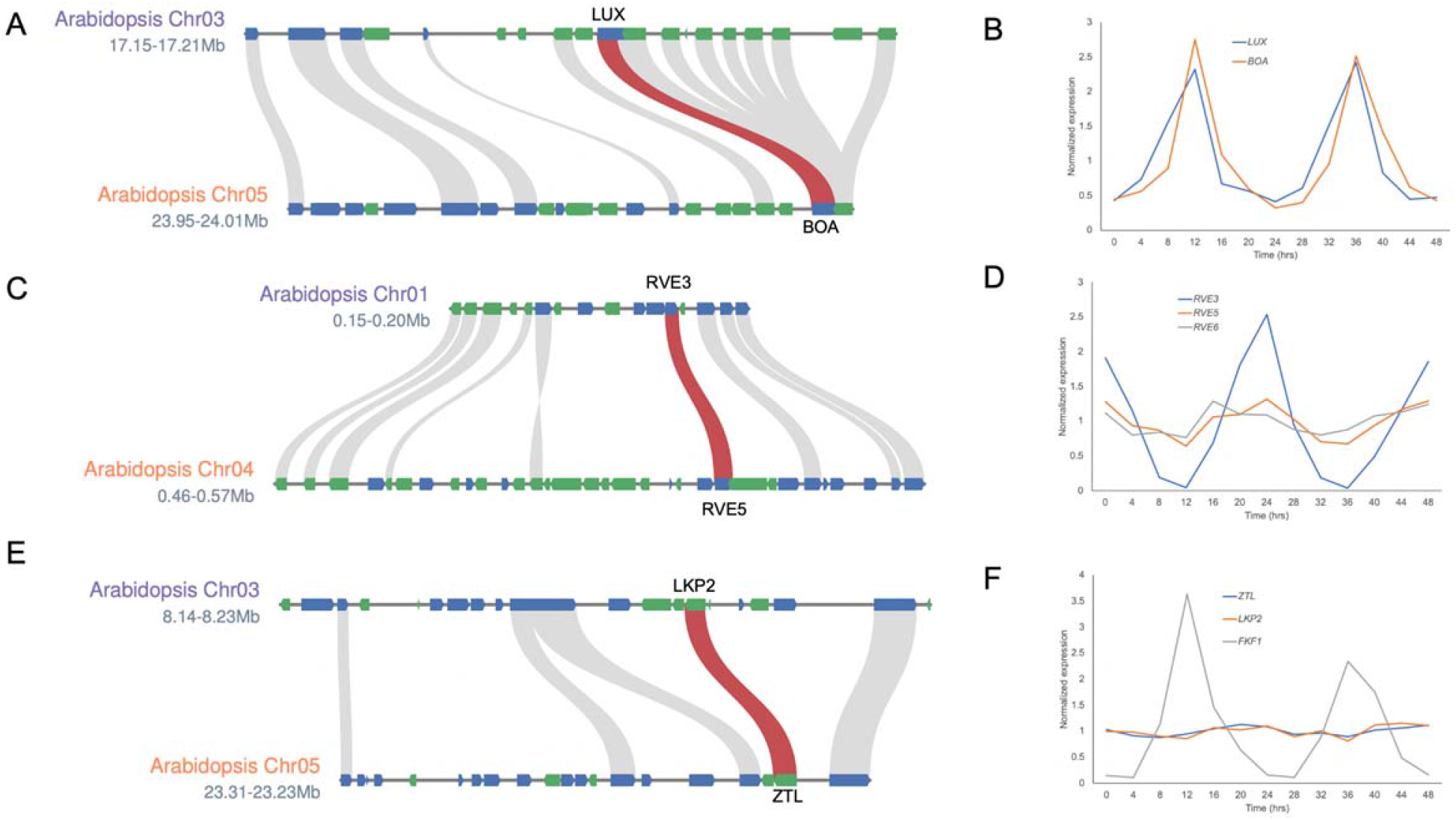
Syntenic orthologs of core circadian clock genes. A) The syntenic block and expression for *LUX* and *BOA*; C-D) The syntenic block and expression for *RVE3* and *RVE5*; and E) The syntenic block and expression for *LKP2* and *ZTL*. Key syntenic relationships (red) and other syntenic genes (grey) for the entire syntenic block. Genes on the positive strand (blue) and negative strand (green).

**Supplemental Figure S3.**
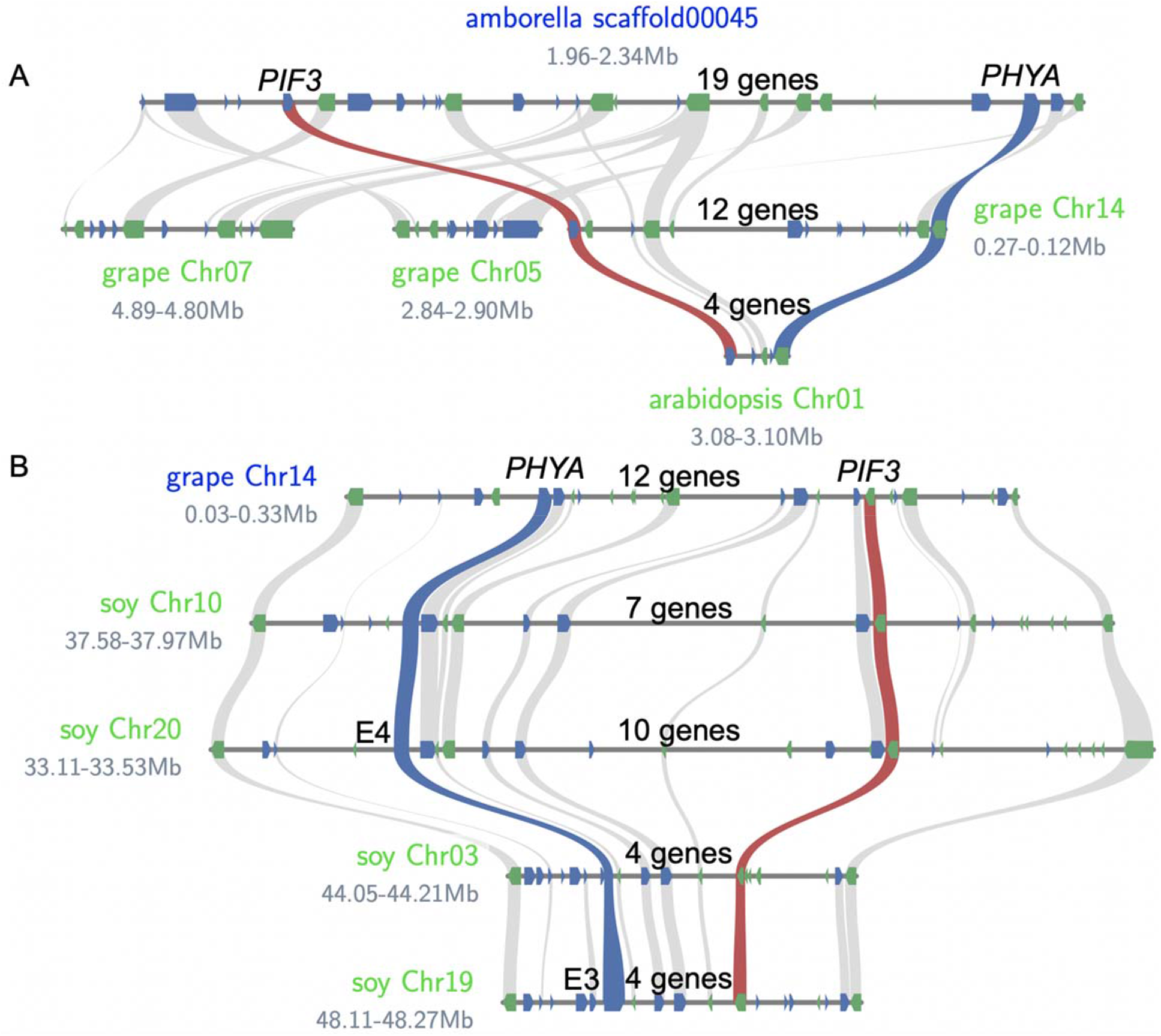
*PIF3* and *PHYA* linkage conserved back to *Amborella*. *PIF3* syntenic relationships (red), *PHYA* syntenic relationships (blue) and other syntenic genes (grey) for the entire syntenic block. Genes on the positive strand (blue) and negative strand (green). A) Syntenic blocks between *Amborella* (*Amborella trichopoda*), grape (*Vitis vinifera*) and *Arabidopsis* (*Arabidopsis thaliana*). B) Syntenic blocks between grape (*Vitis vinifera*) and soy (Glycine max).

**Supplemental Figure 4.**
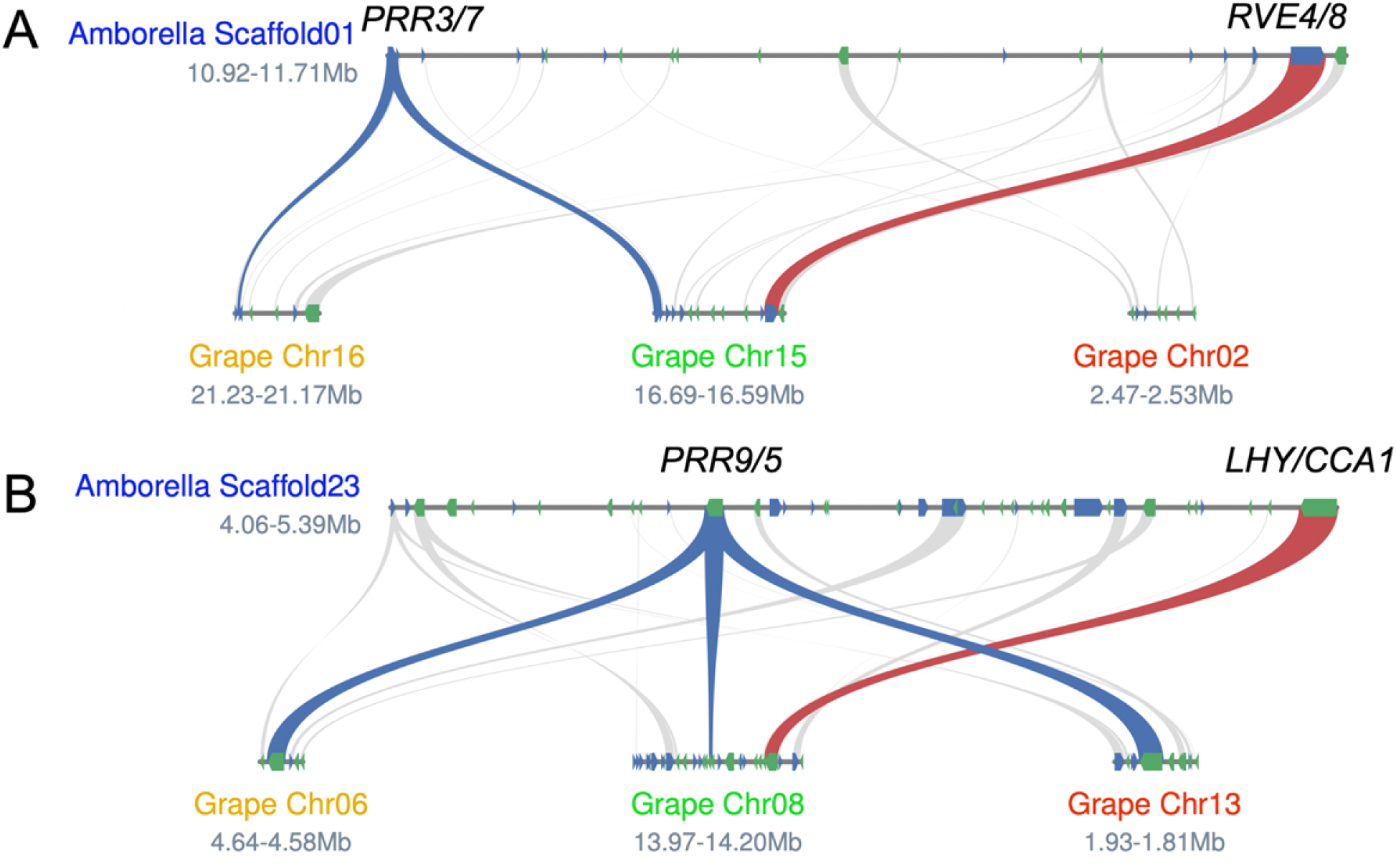
Syntenic sMYB-PRR pairs between *Amborella* and Grape. *LHY/CCA1* syntenic relationships (red), *PRR3/5/7/9* syntenic relationships (blue) and other syntenic genes (grey) for the entire syntenic block. Genes on the positive strand (blue) and negative strand (green). A) Syntenic blocks between *Amborella* (*Amborella trichopoda*) and grape (*Vitis vinifera*) for the *RVE4/8-PRR3/7*. B) A) Syntenic blocks between *Amborella* (*Amborella trichopoda*) and grape (*Vitis vinifera*) for the *LHY/CCA1-PRR5/9*.

**Supplemental Figure S5.**
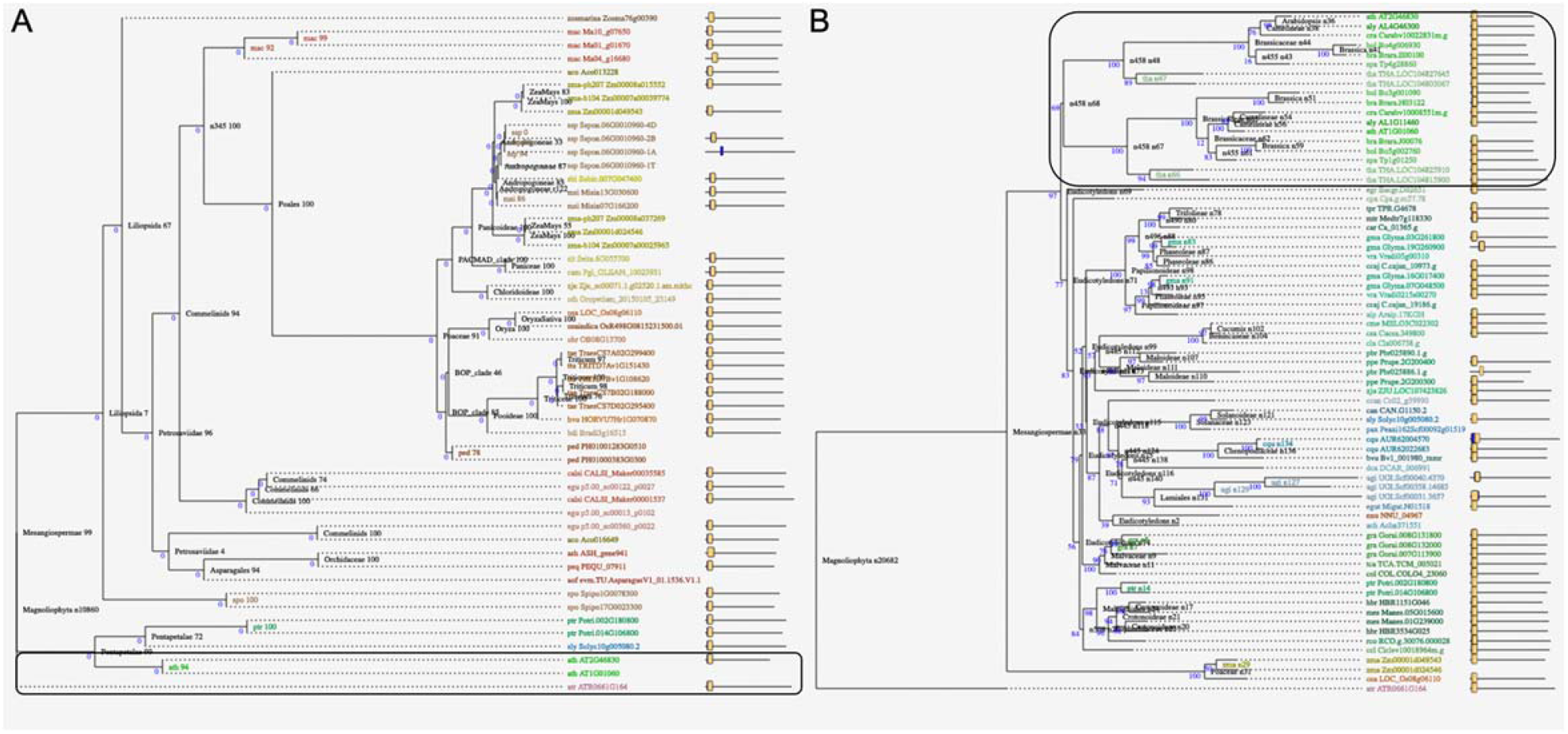
*CCA1/LHY* lineage across monocots and eudicots. Both trees are pre-generated from the PLAZA dicot and monocot web pages. Gene names are on the tips of the tree and domain structure is depicted to the left. A) Monocot *LHY/CCA1* tree; and B) Eudicot *LHY/CCA1* tree. *CCA1* genes boxed.

**Supplemental Figure S6.**
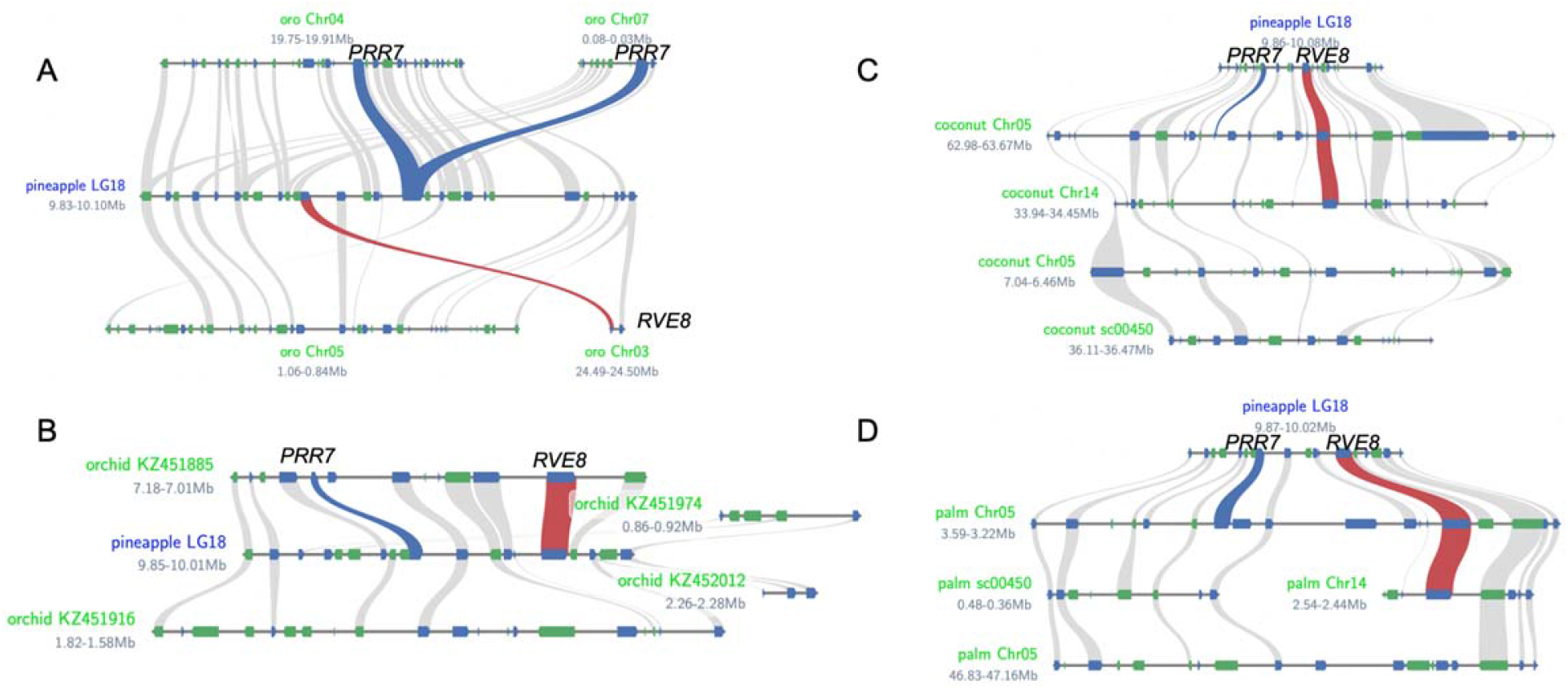
The *sMYB-PRR* syntenic block in pineapple reveal relationships across monocots. *LHY/CCA1/RVE* syntenic relationships (red), *PRR3/5/7/9* syntenic relationships (blue) and other syntenic genes (grey) for the entire syntenic block. Genes on the positive strand (blue) and negative strand (green). A) Pineapple (*Ananas comosus*) versus oro (*Oropetium thomaeum*); B) pineapple versus orchid (*Apostasia shenzhenica*); C) pineapple versus coconut (*Cocos nucifera*); D) pineapple versus palm (*Elaeis guineensis*).

**Supplemental Figure S7.**
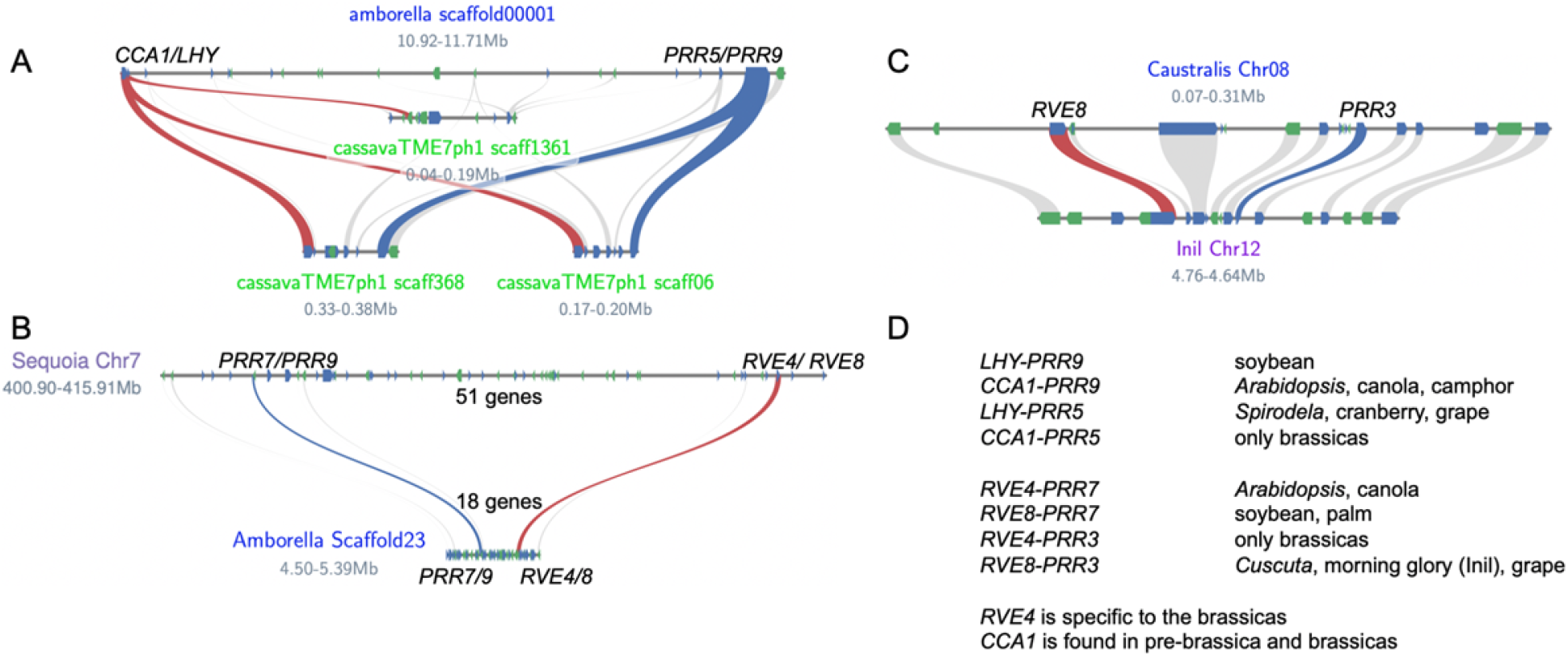
*LHY/CCA1/RVE* syntenic relationships (red), *PRR3/5/7/9* syntenic relationships (blue) and other syntenic genes (grey) for the entire syntenic block. Genes on the positive strand (blue) and negative strand (green). A) *Amborella* versus cassava (*Manihot esculenta*); B) Sequoia (*Sequoiadendron giganteum*) versus *Amborella*; C) Cuscuta (Caustralis; *Cuscuta australis*) versus Inil (*Ipomoea nil*) D) Eight different *sMYB-PRR* combinations and genome examples for each; this is not meant to be an exhaustive list.

**Supplemental Figure S8.**
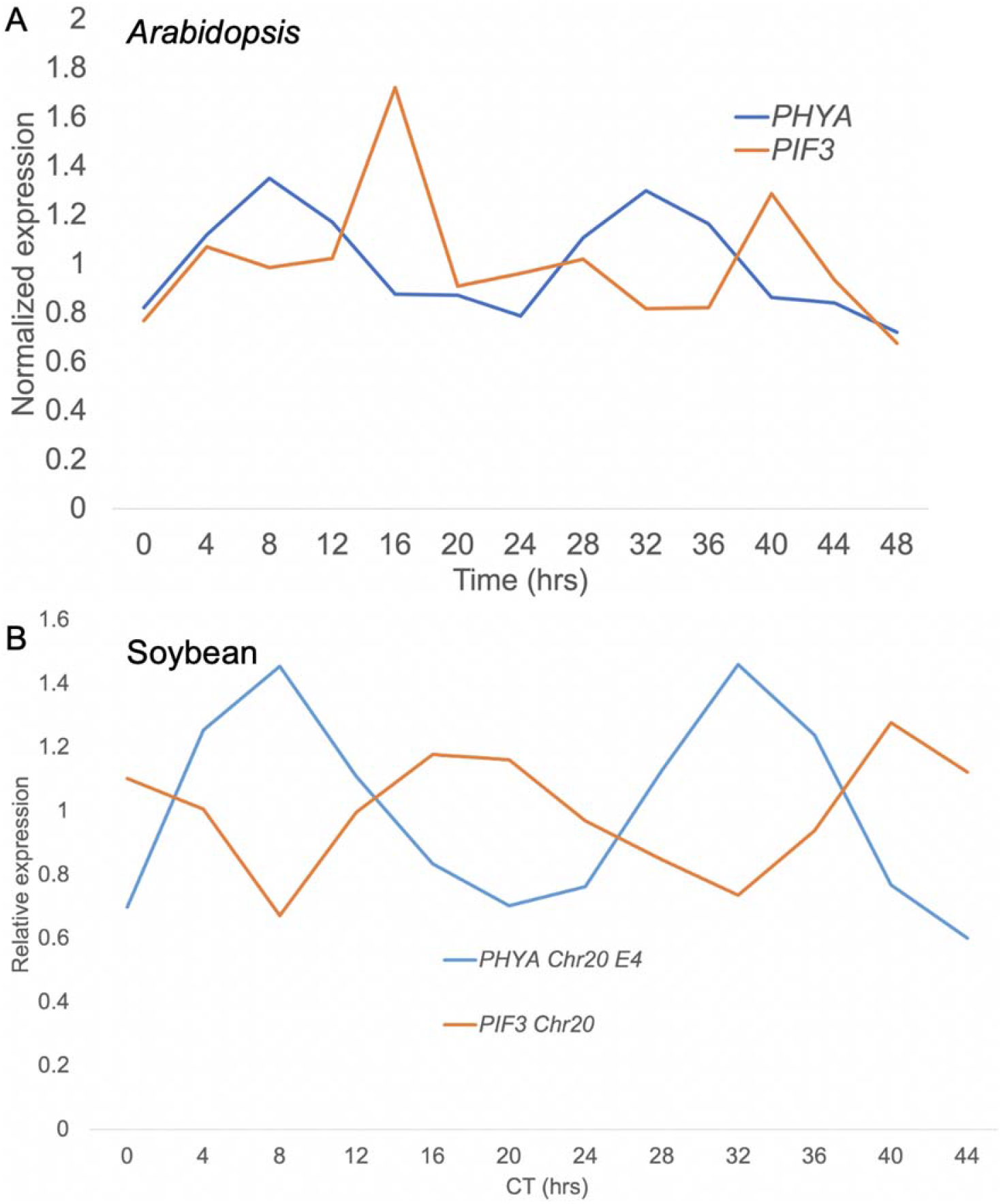
*PHYA-PIF3* are expressed at distinct times of day in *Arabidopsis* and soybean. A) *Arabidopsis PHYA* (blue) shows peak expression at ZT11, while *PIF3* (orange) has peak expression ZT16. B) One syntenic pair of the four *PHYA-PIF3* linkages in soybean robustly cycles under circadian conditions with *PHYA* (blue) peaking at CT8 and *PIF3* (orange) peaking at CT17. ZT; Zeitgeber Time. CT; Circadian Time.

**Supplemental Figure S9.**
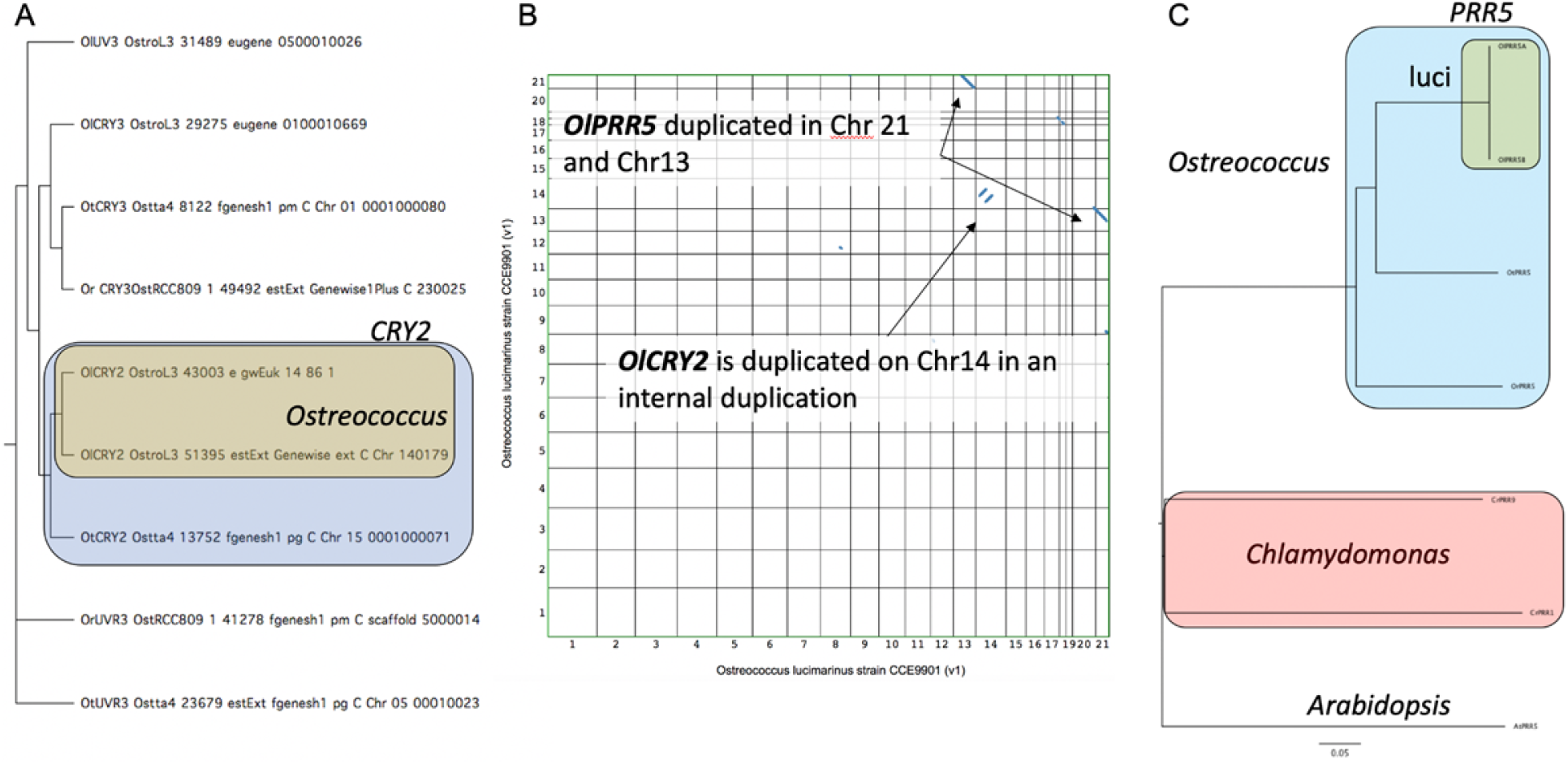
Circadian clock and light signalling genes are duplicated in Ostreococcus. A) *CRYPTOCHROME* (*CRY/UVR*) family in Ostreococcus; all CRY2 genes (blue box) and duplicated *CRY2* genes in *O. lucimarinus* (brown box). B) Dotplot of *O. lucimarinus* (Ol) showing both the *CRY2* and *PRR5* duplications. C) *PRR* genes across *Arabidopsis*, *Chlamydomonas* (red box) and *Ostreococcus* (blue box). All of the *PRR* from *Ostreococcus* (*Ol*, *O. lucimarinus*; *Or*, *O. tauri*; and *Or*) and the duplicated *PRR5* in Ol (luci).

### Supplemental Tables

**Supplemental Table S1.** Arabidopsis circadian clock, light signalling and flowering time genes.

|  | sMYB |  |  | light/flowering |
| --- | --- | --- | --- | --- |
| CCA1 | AT2G46830 |  | ARR3 | At1g59940 |
| LHY | AT1G01060 |  | ARR4 | At1g10470 |
| RVE1 | AT5G17300 |  | bHLH69 | At4g30980 |
| RVE2 | AT5G37260 |  | bHLH92 | At5g43650 |
| RVE3 | AT1G01520 |  | CCR1 | At4g39260 |
| RVE4 | AT5G02840 |  | CCR2 | At2g21660 |
| RVE5 | AT4G01280 |  | COL2 | At3g02380 |
| RVE6 | AT5G52660 |  | COL9 | At3g07650 |
| RVE7 | AT1G18330 |  | COP1 | At2g32950 |
| RVE8 | AT3G09600 |  | CRB | At1g09340 |
|  |  |  | CRY1 | At4g08920 |
|  | PRR |  | CRY2 | At1g04400 |
| PRR1 | AT5G61380 |  | DET1 | At4g10180 |
| PRR3 | AT5G60100 |  | EID1 | At4g02440 |
| PRR5 | AT5G24470 |  | FHY3 | At3g22170 |
| PRR7 | AT5G02810 |  | FIO1 | At2g21070 |
| PRR9 | AT2G46790 |  | FLC | At5g10140 |
|  |  |  | FT | At1g65480 |
|  | ZTL |  | HYH | At3g17609 |
| ZTL | AT5G57360 |  | LIP1 | At2g20860 |
| LKP2 | AT2G18915 |  | LNK1 | AT5G64170 |
| FKF1 | AT1G68050 |  | LNK2 | AT3G54500 |
|  |  |  | PHYA | At1g09570 |
| GI | AT1G22770 |  | PHYB | At2g18790 |
| ELF3 | AT2G25930 |  | PIF3 | At1g09530 |
| ELF4 | AT2G40080 |  | PRMT5 | At4g31120 |
| PCL1/LUX | AT3G46640 |  | SEC | At3g04240 |
| LUX-like | AT5G59570 |  | SFR6 | At4g04920 |
|  |  |  | SPA1 | At2g46340 |
| TEJ | AT2G31870 |  | SPY | At3g11540 |
| TIC | AT3G22380 |  | SRR1 | At5g59560 |
| CDF3 | AT3G47500 |  | STN7 | At1g68830 |
| CHE | AT5G08330 |  |  |  |
| CKB3 | AT3G60250 |  |  |  |
| CKB4 | AT2G44680 |  |  |  |

**Supplemental Table S2.**
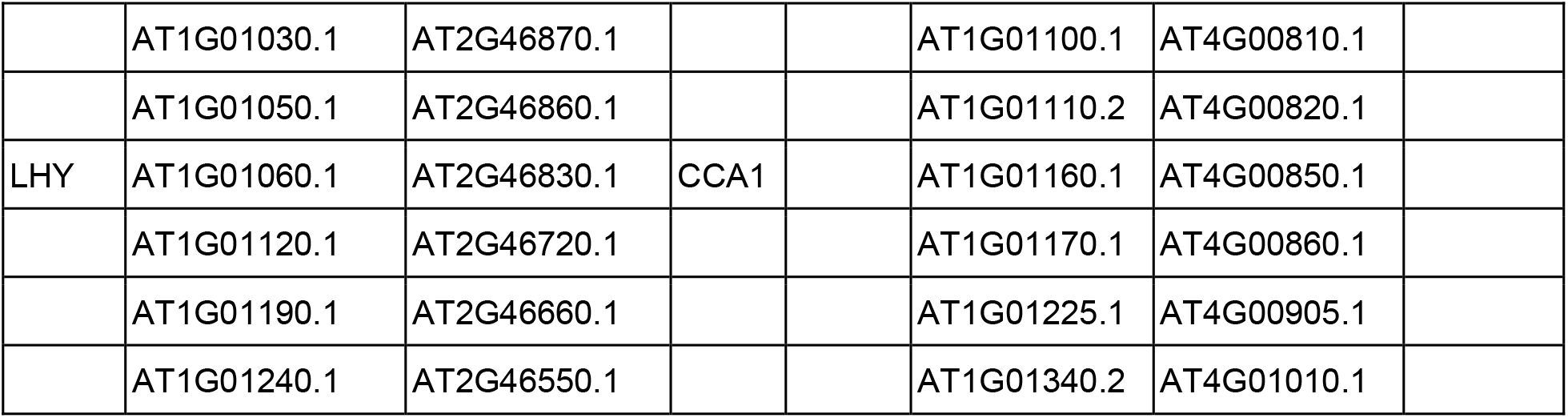
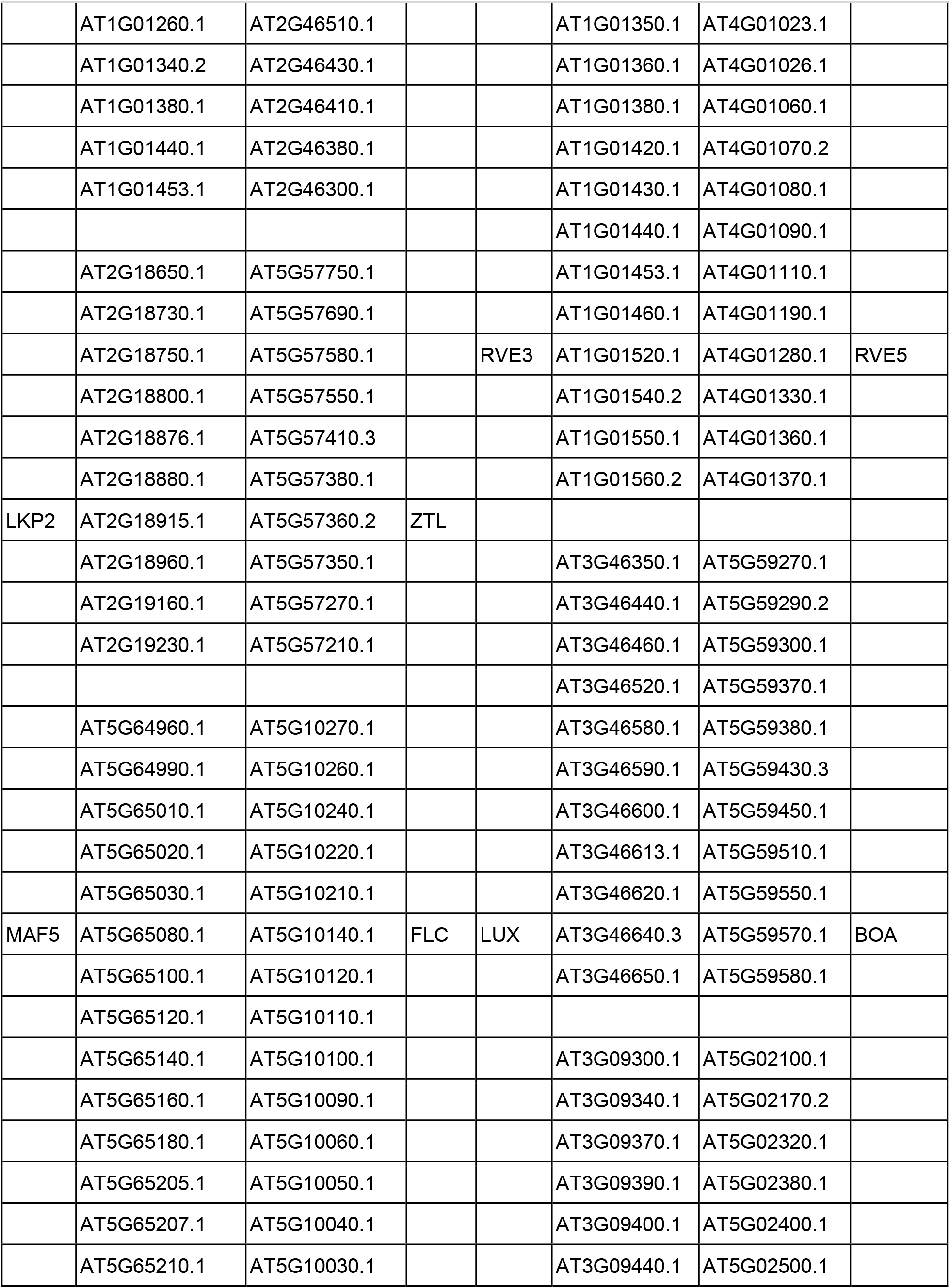
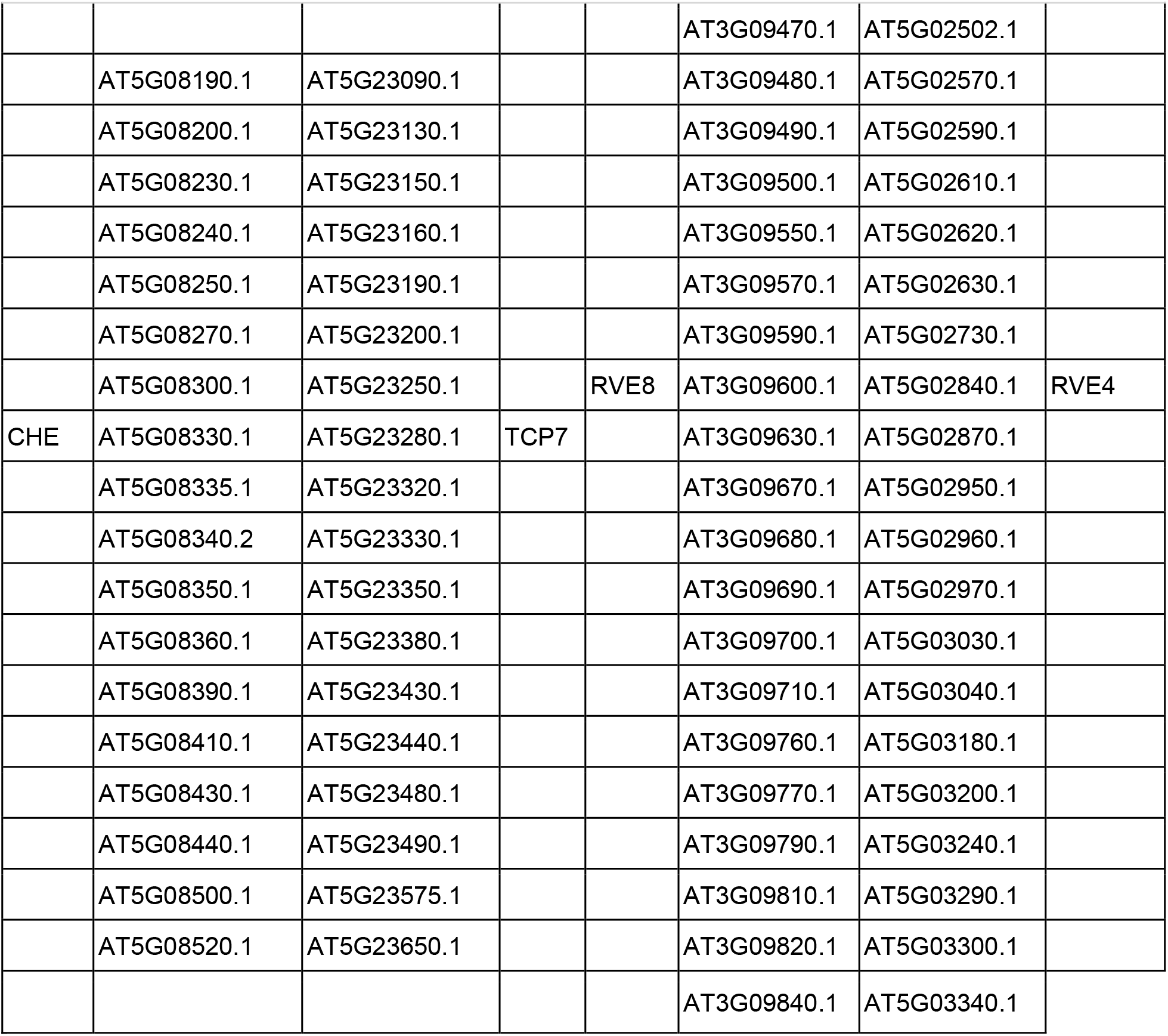
Syntenic regions for core circadian clock genes in *Arabidopsis*.

|  |  |  |  |  |  |  |
| --- | --- | --- | --- | --- | --- | --- |
|  | AT1G01030.1 | AT2G46870.1 |  |  | AT1G01100.1 | AT4G00810.1 |
|  | AT1G01050.1 | AT2G46860.1 |  |  | AT1G01110.2 | AT4G00820.1 |
| LHY | AT1G01060.1 | AT2G46830.1 | CCA1 |  | AT1G01160.1 | AT4G00850.1 |
|  | AT1G01120.1 | AT2G46720.1 |  |  | AT1G01170.1 | AT4G00860.1 |
|  | AT1G01190.1 | AT2G46660.1 |  |  | AT1G01225.1 | AT4G00905.1 |
|  | AT1G01240.1 | AT2G46550.1 |  |  | AT1G01340.2 | AT4G01010.1 |

|  |  |  |  |  |  |  |  |
| --- | --- | --- | --- | --- | --- | --- | --- |
|  | AT1G01260.1 | AT2G46510.1 |  |  | AT1G01350.1 | AT4G01023.1 |  |
|  | AT1G01340.2 | AT2G46430.1 |  |  | AT1G01360.1 | AT4G01026.1 |  |
|  | AT1G01380.1 | AT2G46410.1 |  |  | AT1G01380.1 | AT4G01060.1 |  |
|  | AT1G01440.1 | AT2G46380.1 |  |  | AT1G01420.1 | AT4G01070.2 |  |
|  | AT1G01453.1 | AT2G46300.1 |  |  | AT1G01430.1 | AT4G01080.1 |  |
|  |  |  |  |  | AT1G01440.1 | AT4G01090.1 |  |
|  | AT2G18650.1 | AT5G57750.1 |  |  | AT1G01453.1 | AT4G01110.1 |  |
|  | AT2G18730.1 | AT5G57690.1 |  |  | AT1G01460.1 | AT4G01190.1 |  |
|  | AT2G18750.1 | AT5G57580.1 |  | RVE3 | AT1G01520.1 | AT4G01280.1 | RVE5 |
|  | AT2G18800.1 | AT5G57550.1 |  |  | AT1G01540.2 | AT4G01330.1 |  |
|  | AT2G18876.1 | AT5G57410.3 |  |  | AT1G01550.1 | AT4G01360.1 |  |
|  | AT2G18880.1 | AT5G57380.1 |  |  | AT1G01560.2 | AT4G01370.1 |  |
| LKP2 | AT2G18915.1 | AT5G57360.2 | ZTL |  |  |  |  |
|  | AT2G18960.1 | AT5G57350.1 |  |  | AT3G46350.1 | AT5G59270.1 |  |
|  | AT2G19160.1 | AT5G57270.1 |  |  | AT3G46440.1 | AT5G59290.2 |  |
|  | AT2G19230.1 | AT5G57210.1 |  |  | AT3G46460.1 | AT5G59300.1 |  |
|  |  |  |  |  | AT3G46520.1 | AT5G59370.1 |  |
|  | AT5G64960.1 | AT5G10270.1 |  |  | AT3G46580.1 | AT5G59380.1 |  |
|  | AT5G64990.1 | AT5G10260.1 |  |  | AT3G46590.1 | AT5G59430.3 |  |
|  | AT5G65010.1 | AT5G10240.1 |  |  | AT3G46600.1 | AT5G59450.1 |  |
|  | AT5G65020.1 | AT5G10220.1 |  |  | AT3G46613.1 | AT5G59510.1 |  |
|  | AT5G65030.1 | AT5G10210.1 |  |  | AT3G46620.1 | AT5G59550.1 |  |
| MAF5 | AT5G65080.1 | AT5G10140.1 | FLC | LUX | AT3G46640.3 | AT5G59570.1 | BOA |
|  | AT5G65100.1 | AT5G10120.1 |  |  | AT3G46650.1 | AT5G59580.1 |  |
|  | AT5G65120.1 | AT5G10110.1 |  |  |  |  |  |
|  | AT5G65140.1 | AT5G10100.1 |  |  | AT3G09300.1 | AT5G02100.1 |  |
|  | AT5G65160.1 | AT5G10090.1 |  |  | AT3G09340.1 | AT5G02170.2 |  |
|  | AT5G65180.1 | AT5G10060.1 |  |  | AT3G09370.1 | AT5G02320.1 |  |
|  | AT5G65205.1 | AT5G10050.1 |  |  | AT3G09390.1 | AT5G02380.1 |  |
|  | AT5G65207.1 | AT5G10040.1 |  |  | AT3G09400.1 | AT5G02400.1 |  |
|  | AT5G65210.1 | AT5G10030.1 |  |  | AT3G09440.1 | AT5G02500.1 |  |

|  |  |  |  |  |  |  |  |
| --- | --- | --- | --- | --- | --- | --- | --- |
|  |  |  |  |  | AT3G09470.1 | AT5G02502.1 |  |
|  | AT5G08190.1 | AT5G23090.1 |  |  | AT3G09480.1 | AT5G02570.1 |  |
|  | AT5G08200.1 | AT5G23130.1 |  |  | AT3G09490.1 | AT5G02590.1 |  |
|  | AT5G08230.1 | AT5G23150.1 |  |  | AT3G09500.1 | AT5G02610.1 |  |
|  | AT5G08240.1 | AT5G23160.1 |  |  | AT3G09550.1 | AT5G02620.1 |  |
|  | AT5G08250.1 | AT5G23190.1 |  |  | AT3G09570.1 | AT5G02630.1 |  |
|  | AT5G08270.1 | AT5G23200.1 |  |  | AT3G09590.1 | AT5G02730.1 |  |
|  | AT5G08300.1 | AT5G23250.1 |  | RVE8 | AT3G09600.1 | AT5G02840.1 | RVE4 |
| CHE | AT5G08330.1 | AT5G23280.1 | TCP7 |  | AT3G09630.1 | AT5G02870.1 |  |
|  | AT5G08335.1 | AT5G23320.1 |  |  | AT3G09670.1 | AT5G02950.1 |  |
|  | AT5G08340.2 | AT5G23330.1 |  |  | AT3G09680.1 | AT5G02960.1 |  |
|  | AT5G08350.1 | AT5G23350.1 |  |  | AT3G09690.1 | AT5G02970.1 |  |
|  | AT5G08360.1 | AT5G23380.1 |  |  | AT3G09700.1 | AT5G03030.1 |  |
|  | AT5G08390.1 | AT5G23430.1 |  |  | AT3G09710.1 | AT5G03040.1 |  |
|  | AT5G08410.1 | AT5G23440.1 |  |  | AT3G09760.1 | AT5G03180.1 |  |
|  | AT5G08430.1 | AT5G23480.1 |  |  | AT3G09770.1 | AT5G03200.1 |  |
|  | AT5G08440.1 | AT5G23490.1 |  |  | AT3G09790.1 | AT5G03240.1 |  |
|  | AT5G08500.1 | AT5G23575.1 |  |  | AT3G09810.1 | AT5G03290.1 |  |
|  | AT5G08520.1 | AT5G23650.1 |  |  | AT3G09820.1 | AT5G03300.1 |  |
|  |  |  |  |  | AT3G09840.1 | AT5G03340.1 |  |

**Supplemental Table S3.**
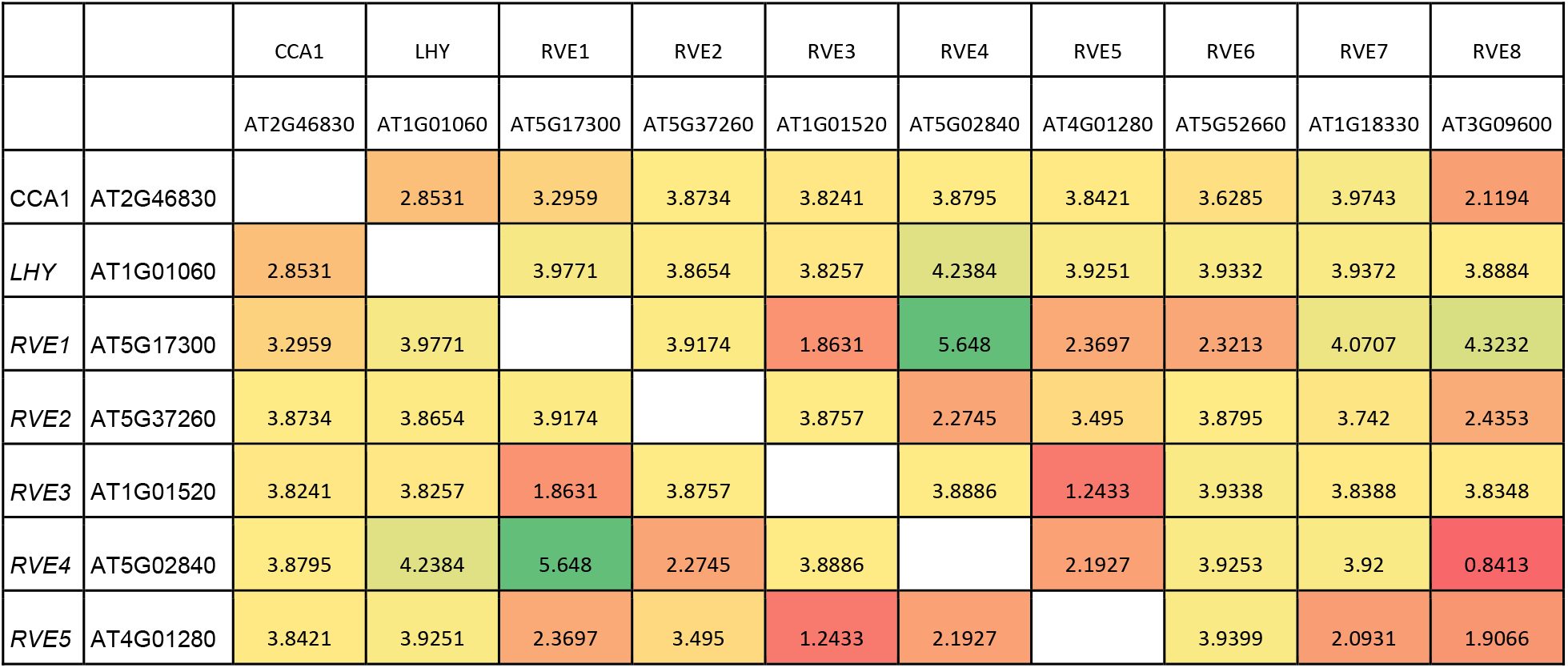
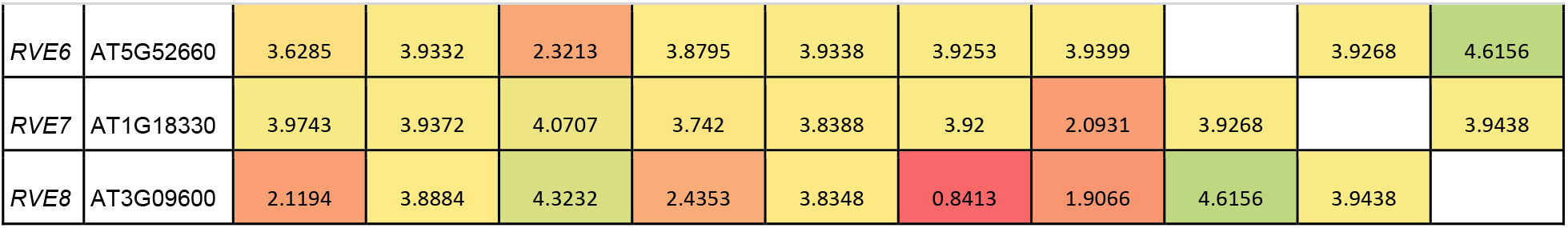
Synonymous substitution (Ks) across *Arabidopsis* sMYB proteins.

|  |  | CCA1 | LHY | RVE1 | RVE2 | RVE3 | RVE4 | RVE5 | RVE6 | RVE7 | RVE8 |
| --- | --- | --- | --- | --- | --- | --- | --- | --- | --- | --- | --- |
|  |  | AT2G46830 | AT1G01060 | AT5G17300 | AT5G37260 | AT1G01520 | AT5G02840 | AT4G01280 | AT5G52660 | AT1G18330 | AT3G09600 |
| CCA1 | AT2G46830 |  | 2.8531 | 3.2959 | 3.8734 | 3.8241 | 3.8795 | 3.8421 | 3.6285 | 3.9743 | 2.1194 |
| LHY | AT1G01060 | 2.8531 |  | 3.9771 | 3.8654 | 3.8257 | 4.2384 | 3.9251 | 3.9332 | 3.9372 | 3.8884 |
| RVE1 | AT5G17300 | 3.2959 | 3.9771 |  | 3.9174 | 1.8631 | 5.648 | 2.3697 | 2.3213 | 4.0707 | 4.3232 |
| RVE2 | AT5G37260 | 3.8734 | 3.8654 | 3.9174 |  | 3.8757 | 2.2745 | 3.495 | 3.8795 | 3.742 | 2.4353 |
| RVE3 | AT1G01520 | 3.8241 | 3.8257 | 1.8631 | 3.8757 |  | 3.8886 | 1.2433 | 3.9338 | 3.8388 | 3.8348 |
| RVE4 | AT5G02840 | 3.8795 | 4.2384 | 5.648 | 2.2745 | 3.8886 |  | 2.1927 | 3.9253 | 3.92 | 0.8413 |
| RVE5 | AT4G01280 | 3.8421 | 3.9251 | 2.3697 | 3.495 | 1.2433 | 2.1927 |  | 3.9399 | 2.0931 | 1.9066 |
| RVE6 | AT5G52660 | 3.6285 | 3.9332 | 2.3213 | 3.8795 | 3.9338 | 3.9253 | 3.9399 |  | 3.9268 | 4.6156 |
| RVE7 | AT1G18330 | 3.9743 | 3.9372 | 4.0707 | 3.742 | 3.8388 | 3.92 | 2.0931 | 3.9268 |  | 3.9438 |
| RVE8 | AT3G09600 | 2.1194 | 3.8884 | 4.3232 | 2.4353 | 3.8348 | 0.8413 | 1.9066 | 4.6156 | 3.9438 |  |

**Supplemental Table S4.** Synonymous substitution (Ks) across *Arabidopsis* PRR proteins.

|  |  | PRR1 | PRR3 | PRR5 | PRR7 | PRR9 |
| --- | --- | --- | --- | --- | --- | --- |
|  |  | AT5G61380 | AT5G60100 | AT5G24470 | AT5G02810 | AT2G46790 |
| PRR1 | AT5G61380 |  | 4.1647 | 4.2759 | 4.3028 | 4.2329 |
| PRR3 | AT5G60100 | 4.1647 |  | 4.2775 | 4.2402 | 2.641 |
| PRR5 | AT5G24470 | 4.2759 | 4.2775 |  | 4.381 | 2.328 |
| PRR7 | AT5G02810 | 4.3028 | 4.2402 | 4.381 |  | 2.5112 |
| PRR9 | AT2G46790 | 4.2329 | 2.641 | 2.328 | 2.5112 |  |

**Supplemental Table S5.** Genetic linkages between *CCA1/LHY-PRR5/9* and *RVE4/8-PRR3/7* from PLAZA dicot 4.5. For each species the number of genetic linkages is presented.

| abbreviation | common_name | tax_id | pubmed | LHY/CCA1-PRR5/9 | RVE4/8-PRR3/7 | total |
| --- | --- | --- | --- | --- | --- | --- |
| ach | *Actinidia chinensis* | 3625 | 24136039 | 0 | 0 | 0 |
| Ahy | *Amaranthus hypochondriacus* | NA | NA | 1 | 2 | 3 |
| aip | *Arachis ipaensis* | 130453 | 26901068 | 0 | 1 | 1 |
| aly | *Arabidopsis lyrata* | 59689 | 26382944 | 1 | 1 | 2 |
| ath | *Arabidopsis thaliana* | 3702 | 27862469 | 1 | 1 | 2 |
| atr | *Amborella trichopoda* | 13333 | 24357323 | 1 | 1 | 2 |
| bol | *Brassica oleracea* | 109376 | 24852848 | 1 | 0 | 1 |
| bra | *Brassica rapa* | 3711 | 21873998 | 0 | 1 | 1 |
| bvu | *Beta vulgaris* | 161934 | 24352233 | 1 | 0 | 1 |
| can | *Capsicum annuum* | 4072 | 24441736 | 0 | 0 | 0 |
| car | *Cicer arietinum* | 3827 | 23354103 | 1 | 1 | 2 |
| ccaj | *Cajanus cajan* | 3821 | 22057054 | 0 | 0 | 0 |
| ccan | *Coffea canephora* | 49390 | 25190796 | 1 | 1 | 2 |
| ccl | *Citrus clementina* | 85681 | 24908277 | 1 | 1 | 2 |
| cla | *Citrullus lanatus* | 3654 | 23179023 | 0 | 0 | 0 |
| cme | *Cucumis melo* | 3656 | 22753475 | 0 | 0 | 0 |
| col | *Corchorus olitorius* | 93759 | 28134914 | 1 | 0 | 1 |
| cpa | *Carica papaya* | 3649 | 18432245 | 0 | 0 | 0 |
| cqu | *Chenopodium quinoa* | 63459 | 28178233 | 2 | 0 | 2 |
| cre | *Chlamydomonas reinhardtii* | 3055 | 17932292 | 0 | 0 | 0 |
| cru | *Capsella rubella* | 81985 | 23749190 | 1 | 1 | 2 |
| csa | *Cucumis sativus L.* | 3659 | NA | 0 | 0 | 0 |
| dca | *Daucus carota* | 4039 | 27158781 | 0 | 0 | 0 |
| egr | *Eucalyptus grandis* | 71139 | 24919147 | 0 | 0 | 0 |
| egut | *Erythranthe guttata* | 4155 | 24225854 | 0 | 1 | 1 |
| fve | *Fragaria vesca* | 57918 | 21186353 | 0 | 1 | 1 |
| gma | *Glycine max* | 3847 | 20075913 | 4 | 2 | 6 |
| gra | *Gossypium raimondii* | 29730 | 22922876 | 1 | 1 | 2 |
| hbr | *Hevea brasiliensis* | 3981 | 27255837 | 1 | 2 | 3 |
| mco | *Micromonas commoda* | 296587 | 19359590 | 0 | 0 | 0 |
| mdo | *Malus domestica* | 3750 | 20802477 | 0 | 0 | 0 |
| mes | *Manihot esculenta* | 3983 | 22523606 | 0 | 0 | 0 |
| mpo | *Marchantia polymorpha* | 3197 | 28985561 | 0 | 0 | 0 |
| mtr | *Medicago truncatula* | 3880 | 22089132 | 1 | 1 | 2 |
| nnu | *Nelumbo nucifera* | 4432 | 23663246 | 0 | 1 | 1 |
| osa | *Oryza sativa ssp. japonica* | 39947 | 16100779 | 0 | 0 | 0 |
| pab | *Picea abies* | 3329 | 23698360 | 0 | 0 | 0 |
| pax | *Petunia axillaris* | 33119 | 27255838 | 0 | 0 | 0 |
| pbr | *Pyrus bretschneideri* | 225117 | 23149293 | 0 | 1 | 1 |
| ppa | *Physcomitrella patens* | 3218 | 18079367 | 0 | 0 | 0 |
| ppe | *Prunus persica* | 3760 | 23525075 | 0 | 1 | 1 |
| ptr | *Populus trichocarpa* | 3694 | 16973872 | 2 | 0 | 2 |
| rco | *Ricinus communis* | 3988 | 20729833 | 0 | 1 | 1 |
| sly | *Solanum lycopersicum* | 4081 | 22660326 | 1 | 0 | 1 |
| smo | *Selaginella moellendorffii* | 88036 | 21551031 | 0 | 0 | 0 |
| spa | *Schrenkiella parvula* | 98039 | 21822265 | 1 | 1 | 2 |
| stu | *Solanum tuberosum* | 4113 | 21743474 | 0 | 1 | 1 |
| tca | *Theobroma cacao* | 3641 | 21186351 | 1 | 1 | 2 |
| tha | *Tarenaya hassleriana* | 28532 | 23983221 | 1 | 2 | 3 |
| tpr | *Trifolium pratense* | 57577 | 26617401 | 1 | 0 | 1 |
| ugi | *Utricularia gibba* | 13748 | 23665961 | 0 | 0 | 0 |
| vra | *Vigna radiata var. radiata* | 157791 | 25384727 | 1 | 1 | 2 |
| vvi | *Vitis vinifera* | 29760 | 17721507 | 1 | 1 | 2 |
| zju | *Ziziphus jujuba* | 326968 | 25350882 | 0 | 0 | 0 |
| zma | *Zea mays* | 4577 | 19965430 | 0 | 0 | 0 |

**Supplemental Table 6.** Number of syntenic blocks and genetic linkages between *CCA1/LHY-PRR5/9*, *RVE4/8-PRR3/7* and PIF3-PHYA. The number under each individual gene pair is the number of syntenic blocks found per species. The number under the genetic linkages represents the number of species sharing genetic linkages with that species.

| species | Clade | Order | CC<br>A1/<br>LHY | RVE<br>4/8 | PR<br>R3/<br>7 | PR<br>R5/<br>9 | PH<br>YA | PIF3 | RVE4/8-PRR3/7 | LHY/CC<br>A1-PRR5/9 | PIF3-PHYA |
| --- | --- | --- | --- | --- | --- | --- | --- | --- | --- | --- | --- |
| *Amborella trichopoda* | Basal-Angiosperm | Amborellales | 1 | 1 | 1 | 2 | 1 | 1 | 23 | 29 | 42 |
| *Nymphaea tetragona* | Basal-Angiosperm | Nymphaeales | 1 | 0 | 2 | 2 | 1 | 1 | 0 | 2 | 41 |
| *Nelumbo nucifera* | Basal-Eudicots | Proteales | 1 | 1 | 2 | 2 | 0 | 2 | 46 | 50 | 0 |
| *Macleaya cordata* | Basal-Eudicots | Ranunculales | 2 | 1 | 1 | 1 | 1 | 1 | 44 | 6 | 72 |
| *Papaver somniferum* | Basal-Eudicots | Ranunculales | 2 | 2 | 1 | 2 | 2 | 2 | 11 | 50 | 108 |
| *Aquilegia coerulea* | Basal-Eudicots | Ranunculales | 1 | 1 | 2 | 0 | 1 | 1 | 24 | 0 | 1 |
| *Vitis vinifera* | Basal-Eudicots | Vitales | 1 | 1 | 4 | 2 | 1 | 2 | 22 | 56 | 72 |
| *Cinnamomum micranthum* | Magnoliids | Magnoliales | 2 | 0 | 2 | 3 | 1 | 2 | 0 | 59 | 64 |
| *Persea americana* | Magnoliids | Magnoliales | 2 | 0 | 2 | 3 | 1 | 2 | 0 | 40 | 0 |
| *Liriodendron chinense* | Magnoliids | Magnoliales | 3 | 0 | 4 | 2 | 1 | 2 | 1 | 35 | 55 |
| *Spirodela polyrhiza* | Monocots | Alismatales | 2 | 1 | 1 | 1 | 2 | 2 | 0 | 2 | 0 |
| *Zostera marina* | Monocots | Alismatales | 0 | 0 | 0 | 0 | 0 | 2 | 0 | 0 | 0 |
| *Elaeis guineensis* | Monocots | Arecales | 1 | 1 | 3 | 0 | 1 | 4 | 50 | 1 | 41 |
| *Phoenix dactylifera* | Monocots | Arecales | 1 | 1 | 2 | 0 | 1 | 2 | 39 | 0 | 0 |
| *Asparagus officinalis* | Monocots | Asparagales | 1 | 1 | 1 | 0 | 1 | 5 | 17 | 0 | 31 |
| *Apostasia shenzhenica* | Monocots | Asparagales | 1 | 1 | 2 | 0 | 1 | 1 | 6 | 0 | 0 |
| *Phalaenopsis equestris* | Monocots | Asparagales | 1 | 0 | 1 | 0 | 1 | 0 | 0 | 0 | 0 |
| *Xerophyta viscosa* | Monocots | Pandanales | 0 | 1 | 3 | 1 | 1 | 3 | 0 | 0 | 2 |
| *Ananas comosus* | Monocots | Poales | 1 | 1 | 2 | 0 | 1 | 2 | 47 | 0 | 59 |
| *Brachypodium distachyon* | Monocots | Poales | 0 | 0 | 2 | 0 | 0 | 2 | 0 | 0 | 0 |
| *Echinochloa crus-galli* | Monocots | Poales | 0 | 0 | 4 | 0 | 0 | 6 | 0 | 0 | 0 |
| *Hordeum vulgare* | Monocots | Poales | 0 | 0 | 2 | 0 | 0 | 1 | 0 | 0 | 0 |
| *leersia perrieri* | Monocots | Poales | 0 | 0 | 3 | 0 | 0 | 2 | 0 | 0 | 0 |
| *Oropetium thomaeum* | Monocots | Poales | 0 | 0 | 3 | 0 | 0 | 2 | 0 | 0 | 0 |
| *Oryza glaberrima* | Monocots | Poales | 0 | 0 | 2 | 0 | 0 | 1 | 0 | 0 | 0 |
| *Oryza punctata* | Monocots | Poales | 0 | 0 | 3 | 0 | 0 | 2 | 0 | 0 | 0 |
| *Oryza rufipogon* | Monocots | Poales | 0 | 0 | 3 | 0 | 0 | 2 | 0 | 0 | 0 |
| *Oryza sativa* | Monocots | Poales | 0 | 0 | 2 | 0 | 0 | 1 | 0 | 0 | 0 |
| *Saccharum officinarum* | Monocots | Poales | 0 | 0 | 3 | 1 | 0 | 1 | 0 | 0 | 0 |
| *Setaria italica* | Monocots | Poales | 0 | 0 | 2 | 0 | 0 | 2 | 0 | 0 | 0 |
| *Setaria viridis* | Monocots | Poales | 0 | 0 | 2 | 0 | 0 | 2 | 0 | 0 | 0 |
| *Sorghum bicolor* | Monocots | Poales | 0 | 0 | 1 | 0 | 0 | 2 | 0 | 0 | 0 |
| *Triticum turgidum* | Monocots | Poales | 0 | 0 | 4 | 0 | 0 | 2 | 0 | 0 | 0 |
| *Zea mays* | Monocots | Poales | 0 | 0 | 3 | 0 | 1 | 3 | 0 | 0 | 0 |
| *Musa acuminata* | Monocots | Zingiberales | 0 | 1 | 4 | 0 | 2 | 2 | 0 | 0 | 0 |
| *Daucus carota* | Super-Asterids | Apiales | 1 | 1 | 3 | 3 | 2 | 1 | 0 | 0 | 71 |
| *Helianthus annuus* | Super-Asterids | Asterales | 0 | 1 | 7 | 1 | 0 | 0 | 0 | 0 | 0 |
| *Lactuca sativa* | Super-Asterids | Asterales | 1 | 1 | 3 | 1 | 1 | 0 | 53 | 0 | 0 |
| *Amaranthus hypochondriacus* | Super-Asterids | Caryophyllales | 2 | 1 | 2 | 2 | 2 | 2 | 46 | 58 | 0 |
| *Beta vulgaris* | Super-Asterids | Caryophyllales | 1 | 1 | 1 | 2 | 1 | 1 | 0 | 56 | 0 |
| *Chenopodium quinoa* | Super-Asterids | Caryophyllales | 2 | 1 | 3 | 4 | 2 | 2 | 7 | 59 | 0 |
| *Actinidia chinensis* | Super-Asterids | Ericales | 1 | 2 | 5 | 3 | 3 | 3 | 0 | 0 | 131 |
| *Actinidia eriantha* | Super-Asterids | Ericales | 2 | 3 | 6 | 3 | 3 | 4 | 1 | 1 | 85 |
| *Coffea canephora* | Super-Asterids | Gentianales | 1 | 1 | 2 | 1 | 1 | 1 | 27 | 0 | 0 |
| *Olea europaea* | Super-Asterids | Lamiales | 1 | 2 | 4 | 3 | 1 | 3 | 0 | 0 | 0 |
| *Sesamum indicum* | Super-Asterids | Lamiales | 1 | 1 | 1 | 2 | 2 | 1 | 55 | 0 | 76 |
| *Mimulus guttatus* | Super-Asterids | Lamiales | 1 | 1 | 2 | 2 | 2 | 1 | 53 | 0 | 74 |
| *Malania oleifera* | Super-Asterids | Santalales | 1 | 0 | 1 | 1 | 1 | 1 | 0 | 0 | 70 |
| *Kalanchoe fedtschenkoi* | Super-Asterids | Saxifragales | 2 | 1 | 5 | 2 | 2 | 2 | 0 | 17 | 71 |
| *Cuscuta campestris* | Super-Asterids | Solanales | 1 | 2 | 8 | 2 | 2 | 0 | 52 | 0 | 0 |
| *Ipomoea nil* | Super-Asterids | Solanales | 0 | 1 | 3 | 3 | 1 | 2 | 41 | 0 | 0 |
| *Capsicum annuum* | Super-Asterids | Solanales | 1 | 1 | 4 | 2 | 0 | 1 | 1 | 33 | 0 |
| *Capsicum baccatum* | Super-Asterids | Solanales | 2 | 1 | 2 | 2 | 0 | 1 | 0 | 32 | 0 |
| *Capsicum chinense* | Super-Asterids | Solanales | 1 | 1 | 2 | 2 | 0 | 1 | 0 | 0 | 0 |
| *Petunia axillaris* | Super-Asterids | Solanales | 0 | 1 | 3 | 3 | 0 | 2 | 3 | 0 | 0 |
| *Solanum lycopersicum* | Super-Asterids | Solanales | 1 | 1 | 3 | 2 | 0 | 1 | 10 | 34 | 0 |
| *Solanum pennellii* | Super-Asterids | Solanales | 1 | 1 | 2 | 2 | 0 | 1 | 1 | 35 | 0 |
| *Solanum tuberosum* | Super-Asterids | Solanales | 1 | 1 | 0 | 2 | 0 | 1 | 0 | 30 | 0 |
| *Aethionema arabicum* | Super-Rosids | Brassicales | 2 | 2 | 2 | 2 | 1 | 3 | 46 | 51 | 0 |
| *Arabidopsis lyrata* | Super-Rosids | Brassicales | 2 | 2 | 4 | 2 | 1 | 6 | 49 | 50 | 67 |
| *Arabidopsis thaliana* | Super-Rosids | Brassicales | 2 | 2 | 2 | 3 | 1 | 4 | 51 | 52 | 67 |
| *Arabis alpina* | Super-Rosids | Brassicales | 0 | 2 | 1 | 0 | 0 | 5 | 0 | 0 | 1 |
| *Boechera stricta* | Super-Rosids | Brassicales | 2 | 2 | 2 | 2 | 1 | 4 | 45 | 50 | 66 |
| *Brassica napus* | Super-Rosids | Brassicales | 3 | 10 | 7 | 9 | 4 | 15 | 50 | 48 | 61 |
| *Brassica oleracea* | Super-Rosids | Brassicales | 4 | 5 | 3 | 4 | 2 | 8 | 45 | 50 | 84 |
| *Brassica rapa* | Super-Rosids | Brassicales | 4 | 4 | 3 | 3 | 2 | 8 | 42 | 0 | 100 |
| *Camelina sativa* | Super-Rosids | Brassicales | 5 | 8 | 6 | 6 | 3 | 10 | 57 | 51 | 194 |
| *Capsella rubella* | Super-Rosids | Brassicales | 2 | 2 | 3 | 2 | 1 | 4 | 51 | 50 | 67 |
| *Lepidium meyenii* | Super-Rosids | Brassicales | 6 | 8 | 9 | 8 | 4 | 12 | 4 | 59 | 254 |
| *Schrenkiella parvula* | Super-Rosids | Brassicales | 1 | 1 | 2 | 2 | 1 | 5 | 0 | 50 | 65 |
| *Thellungiella halophila* | Super-Rosids | Brassicales | 1 | 2 | 5 | 2 | 1 | 3 | 50 | 1 | 66 |
| *Thellungiella salsuginea* | Super-Rosids | Brassicales | 1 | 2 | 2 | 2 | 1 | 3 | 45 | 1 | 66 |
| *Carica papaya* | Super-Rosids | Brassicales | 1 | 1 | 2 | 2 | 1 | 1 | 0 | 0 | 0 |
| *Cleome gynandra* | Super-Rosids | Brassicales | 3 | 1 | 1 | 4 | 0 | 3 | 0 | 50 | 0 |
| *Tarenaya hassleriana* | Super-Rosids | Brassicales | 5 | 2 | 6 | 4 | 2 | 6 | 53 | 25 | 62 |
| *Begonia fuchsioides* | Super-Rosids | Cucurbitales | 7 | 1 | 2 | 7 | 2 | 2 | 0 | 30 | 0 |
| *Citrullus lanatus* | Super-Rosids | Cucurbitales | 1 | 1 | 1 | 1 | 0 | 1 | 0 | 0 | 0 |
| *Cucumis melo* | Super-Rosids | Cucurbitales | 1 | 1 | 1 | 1 | 0 | 1 | 0 | 1 | 0 |
| *Cucumis sativus* | Super-Rosids | Cucurbitales | 1 | 1 | 1 | 1 | 0 | 1 | 0 | 0 | 0 |
| *Cucurbita maxima* | Super-Rosids | Cucurbitales | 2 | 0 | 2 | 3 | 0 | 2 | 0 | 0 | 0 |
| *Datisca glomerata* | Super-Rosids | Cucurbitales | 1 | 1 | 2 | 1 | 0 | 1 | 50 | 56 | 0 |
| *Ammopiptanthus nanus* | Super-Rosids | Fabales | 0 | 1 | 2 | 3 | 1 | 2 | 50 | 1 | 25 |
| *Arachis duranensis* | Super-Rosids | Fabales | 2 | 2 | 2 | 2 | 3 | 4 | 0 | 59 | 166 |
| *Cajanus cajan* | Super-Rosids | Fabales | 1 | 1 | 1 | 2 | 2 | 4 | 0 | 57 | 148 |
| *Cicer arietinum* | Super-Rosids | Fabales | 1 | 0 | 2 | 2 | 2 | 2 | 0 | 55 | 93 |
| *Glycine max* | Super-Rosids | Fabales | 4 | 4 | 4 | 6 | 4 | 8 | 53 | 61 | 337 |
| *Lotus japonicus* | Super-Rosids | Fabales | 1 | 1 | 1 | 8 | 1 | 3 | 51 | 52 | 75 |
| *Lupinus angustifolius* | Super-Rosids | Fabales | 3 | 1 | 2 | 3 | 3 | 7 | 0 | 55 | 181 |
| *Medicago truncatula* | Super-Rosids | Fabales | 1 | 1 | 2 | 3 | 1 | 3 | 55 | 51 | 92 |
| *Phaseolus vulgaris* | Super-Rosids | Fabales | 3 | 2 | 3 | 3 | 2 | 3 | 53 | 62 | 117 |
| *Trifolium pratense* | Super-Rosids | Fabales | 1 | 1 | 1 | 2 | 1 | 3 | 0 | 54 | 53 |
| *Vigna angularis* | Super-Rosids | Fabales | 2 | 2 | 3 | 3 | 2 | 4 | 31 | 51 | 180 |
| *Vigna radiata* | Super-Rosids | Fabales | 2 | 2 | 4 | 2 | 2 | 3 | 40 | 47 | 165 |
| *Betula pendula* | Super-Rosids | Fagales | 1 | 1 | 2 | 3 | 1 | 2 | 0 | 56 | 22 |
| *Casuarina glauca* | Super-Rosids | Fagales | 1 | 1 | 2 | 2 | 1 | 2 | 48 | 57 | 57 |
| *Quercus robur* | Super-Rosids | Fagales | 2 | 1 | 3 | 1 | 1 | 3 | 52 | 5 | 7 |
| *Carya illinoensis* | Super-Rosids | Fagales | 1 | 0 | 3 | 2 | 0 | 3 | 0 | 0 | 0 |
| *Manihot esculenta* | Super-Rosids | Malpighiales | 2 | 1 | 5 | 3 | 1 | 3 | 16 | 58 | 78 |
| *Ricinus communis* | Super-Rosids | Malpighiales | 1 | 1 | 4 | 2 | 1 | 2 | 54 | 53 | 76 |
| *Linum usitatissimum* | Super-Rosids | Malpighiales | 5 | 2 | 2 | 6 | 0 | 2 | 50 | 7 | 0 |
| *Populus trichocarpa* | Super-Rosids | Malpighiales | 4 | 2 | 2 | 4 | 1 | 3 | 0 | 60 | 77 |
| *Durio zibethinus* | Super-Rosids | Malvales | 2 | 2 | 5 | 4 | 0 | 4 | 57 | 0 | 0 |
| *Gossypium barbadense* | Super-Rosids | Malvales | 5 | 2 | 8 | 9 | 0 | 6 | 47 | 58 | 2 |
| *Gossypium hirsutum* | Super-Rosids | Malvales | 6 | 2 | 8 | 8 | 0 | 6 | 47 | 58 | 2 |
| *Gossypium raimondii* | Super-Rosids | Malvales | 3 | 1 | 5 | 4 | 0 | 3 | 46 | 59 | 0 |
| *Theobroma cacao* | Super-Rosids | Malvales | 1 | 1 | 3 | 2 | 1 | 2 | 52 | 58 | 78 |
| *Punica granatum* | Super-Rosids | Myrtales | 1 | 1 | 2 | 2 | 2 | 3 | 49 | 0 | 74 |
| *Eucalyptus grandis* | Super-Rosids | Myrtales | 1 | 1 | 2 | 1 | 1 | 3 | 0 | 0 | 3 |
| *Trema orientale* | Super-Rosids | Rosales | 1 | 1 | 3 | 2 | 1 | 2 | 51 | 54 | 71 |
| *Morus notabilis* | Super-Rosids | Rosales | 1 | 1 | 5 | 2 | 1 | 2 | 50 | 55 | 75 |
| *Ziziphus jujuba* | Super-Rosids | Rosales | 1 | 0 | 3 | 2 | 3 | 2 | 0 | 2 | 79 |
| *Dryas drummondii* | Super-Rosids | Rosales | 1 | 1 | 2 | 2 | 1 | 1 | 53 | 51 | 75 |
| *Fragaria vesca* | Super-Rosids | Rosales | 2 | 1 | 1 | 2 | 1 | 1 | 51 | 60 | 61 |
| *Malus domestica* | Super-Rosids | Rosales | 2 | 1 | 5 | 3 | 2 | 3 | 57 | 54 | 149 |
| *Prunus mume* | Super-Rosids | Rosales | 1 | 1 | 1 | 1 | 1 | 1 | 53 | 0 | 74 |
| *Prunus persica* | Super-Rosids | Rosales | 2 | 1 | 1 | 2 | 1 | 1 | 52 | 55 | 76 |
| *Pyrus x bretschneideri* | Super-Rosids | Rosales | 2 | 1 | 4 | 2 | 2 | 3 | 54 | 1 | 74 |
| *Rosa chinensis* | Super-Rosids | Rosales | 1 | 1 | 2 | 2 | 1 | 2 | 50 | 58 | 73 |
| *Rubus occidentalis* | Super-Rosids | Rosales | 1 | 1 | 2 | 2 | 1 | 1 | 53 | 1 | 76 |
| *Parasponia andersonii* | Super-Rosids | Rosales | 1 | 1 | 3 | 2 | 1 | 2 | 0 | 32 | 75 |
| *Citrus maxima* | Super-Rosids | Sapindales | 1 | 0 | 3 | 2 | 1 | 2 | 0 | 58 | 58 |
| *Citrus sinensis* | Super-Rosids | Sapindales | 1 | 1 | 5 | 1 | 1 | 2 | 43 | 59 | 52 |
| *Xanthoceras sorbifolium* | Super-Rosids | Sapindales | 1 | 0 | 2 | 2 | 1 | 2 | 0 | 58 | 75 |

**Supplemental Table S7.** Summary of syntenic blocks across 123 plant genome assemblies for circadian genes.

|  | CCA1/LHY | RVE4/8 | PRR3/7 | PRR5/9 | PHYA | PIF3 |
| --- | --- | --- | --- | --- | --- | --- |
| Family name | 3634 | 4022 | 1555 | 1018 | 6934 | 851 |
| Syntenic blocks (#) | 182 | 152 | 345 | 264 | 129 | 330 |
| Most syntenic blocks (#) | 7 | 10 | 9 | 9 | 4 | 15 |
| Fewest syntenic blocks (#) | 0 | 0 | 0 | 0 | 0 | 0 |
| Median syntenic blocks (#) | 1 | 1 | 2 | 2 | 1 | 2 |
| Average syntenic blocks (#) | 1.5 | 1.2 | 2.8 | 2.1 | 1 | 2.7 |
| Zero syntenic blocks | 23 | 28 | 2 | 24 | 37 | 4 |
| Total genomes tested (#) | 123 | 123 | 123 | 123 | 123 | 123 |
| missing (%) | 18.70% | 22.80% | 1.60% | 19.50% | 30.10% | 3.30% |
| present (%) | 81.30% | 77.20% | 98.40% | 80.50% | 69.90% | 96.70% |

